# *A Hierarchical Process Model Links Behavioral Aging and Lifespan in* C. elegans

**DOI:** 10.1101/2021.03.31.437415

**Authors:** Natasha Oswal, Olivier M.F. Martin, Sofia Stroustrup, Monika Anna Matusiak Bruckner, Nicholas Stroustrup

## Abstract

Aging involves a transition from youthful vigor to geriatric infirmity and death. To study the relationship between youthful vigor and lifespan, we developed a new version of “the lifespan machine” that can simultaneously measure age-associated changes in behavior and lifespan at high precision in large populations. Across diverse interventions, we find that behavioral aging and lifespan are not parsimoniously explained as manifestations of a single underlying aging process. Instead, the correlation between youthful vigor and lifespan is better explained as the result of two partially independent aging processes that progress during adulthood under the influence of a shared, systemic factor. Our model provides a framework for separating the direct, targeted effects of lifespan-altering interventions from their broad, systemic effects—supporting future efforts in the rational design of clinical interventions in aging.

## Introduction

Most interventions that extend lifespan—dietary changes^1^, mutations^2 3^, and small molecules^4^—act pleiotropically to effect many organismal phenotypes. Disruption of insulin/IGF signaling, for example, extends lifespan, improves protein homeostasis^5^, increases an individuals’ resistance to heat, redox, and xenobiotic stress, improves pathogen resistance^6^ dramatically extend the period of vigorous, youthful movement^7^, and increases the duration late in life spent in a decrepit state^6–8^. Alterations in mitochondrial metabolism extend lifespan and slow age-associated behavioral declines but also slow development rates^9^, decrease fertility, and increase ROS production^10^. As interventions discovered in model organisms begin to be translated into clinical practice , it becomes crucial to understand whether pleiotropic effects of lifespan-extending interventions represent correctable side-effects or unavoidable, fundamental trade-offs. To do this, we need to understand the causal relationships among the downstream physiological processes influenced by lifespan-extending interventions in aging.

Here, we focus on two macroscopically observable outcomes of aging: behavioral decline and lifespan. Behavioral aging is a complex and high-dimensional process^11^ but we reasoned that the automated microscopy platform “the lifespan machine”^12^ could be applied to simultaneously characterize some age-associated changes in behavior and compare them to lifespan across very large populations. In this way, we could quantify the pleiotropic action of lifespan-extending interventions at high statistical and temporal resolution.

## Results

In most *C. elegans* survival assays, permanent cessation of movement is used as a proxy for death. Though accurate in most contexts, movement cessation is vulnerable to the confounding effects of interventions that reduce individuals’ ability or motivation to move. External stimulation can increase the correlation between movement cessation and death^13^, there may be cases where living animals sill remain alive but unresponsive. To eliminate any ambiguity between behavioral aging and lifespan, we therefore set out to develop a non-behavioral proxy for death that could be automatically identified using our imaging technology.

Previous work^12^ revealed that *C.* elegans exhibit a series of body size changes occurring contemporaneously with the final movement of each individual. Over the last few days of life, individuals exhibited a gradual contraction in apparent body size followed by a rapid expansion that occurred contemporaneously with most individual’s final movement^12^. This “death-associated contraction and expansion” was subsequently validated with alternate imaging approaches, and found in animals killed by acute heat or oxidative stress to involve changes in body wall muscles reminiscent of rigor mortis^14^. Alternative non-behavioral markers of death have been identified^15,16^, but technical challenges currently limit their scalability. So, we set out to develop death-associated expansion as a landmark death in high-throughput automated lifespan assays.

### A classification scheme for behavioral and morphological transitions during nematode aging

Using the lifespan machine, we collected by-hand annotated time-lapse images of 236 wild-type *C. elegans* nematodes housed on agar plates and fed UV-inactivated *E. coli* at 20 °C. In agreement with previous findings^11,12,17,18^, we observed variation in the times at which individuals slowed and cease locomotion late in life^18–20^. We confirmed, that contemporaneous to or shortly after their final movements, individuals exhibit a characteristic rapid decrease and then increase in apparent body size immediately after ceasing movement (Fig. 1a). To facilitate analysis and facilitate comparison to data collected by other methodologies, we devised a staging scheme that divides an individuals’ into distinct behavioral and morphological stages: A “Vigorous movement” period during which animals crawl freely across the plate, a “Weak movement” period during which animals permanently occupy a single plate location but exhibit repetitive head and body postural changes, an “Alive but non-moving” period during which animals exhibit no detectable movement, and an “Expanding” period, during which animals exhibit a characteristic increase in their apparent body size (Fig. 1b). This separation of movement into “vigorous” and “weak” stages hews closely to previous definitions developed for interpreting by-hand assays and used elsewhere^17,18,20^, to which we add one additional stage for living but non-moving individuals. We additionally include two post-death phases: a latent period in which animals show neither movement nor morphological changes, followed by a gradual decrease in apparent body size. According to this classification scheme, we found that wild-type animals lived an average of 25.5±4 days, spending on average 67±1 % moving vigorously, 31±1 % moving weakly, and just 1.5±4% (9 hours) alive but motionless (Fig 1c,d).

**Figure 1:**
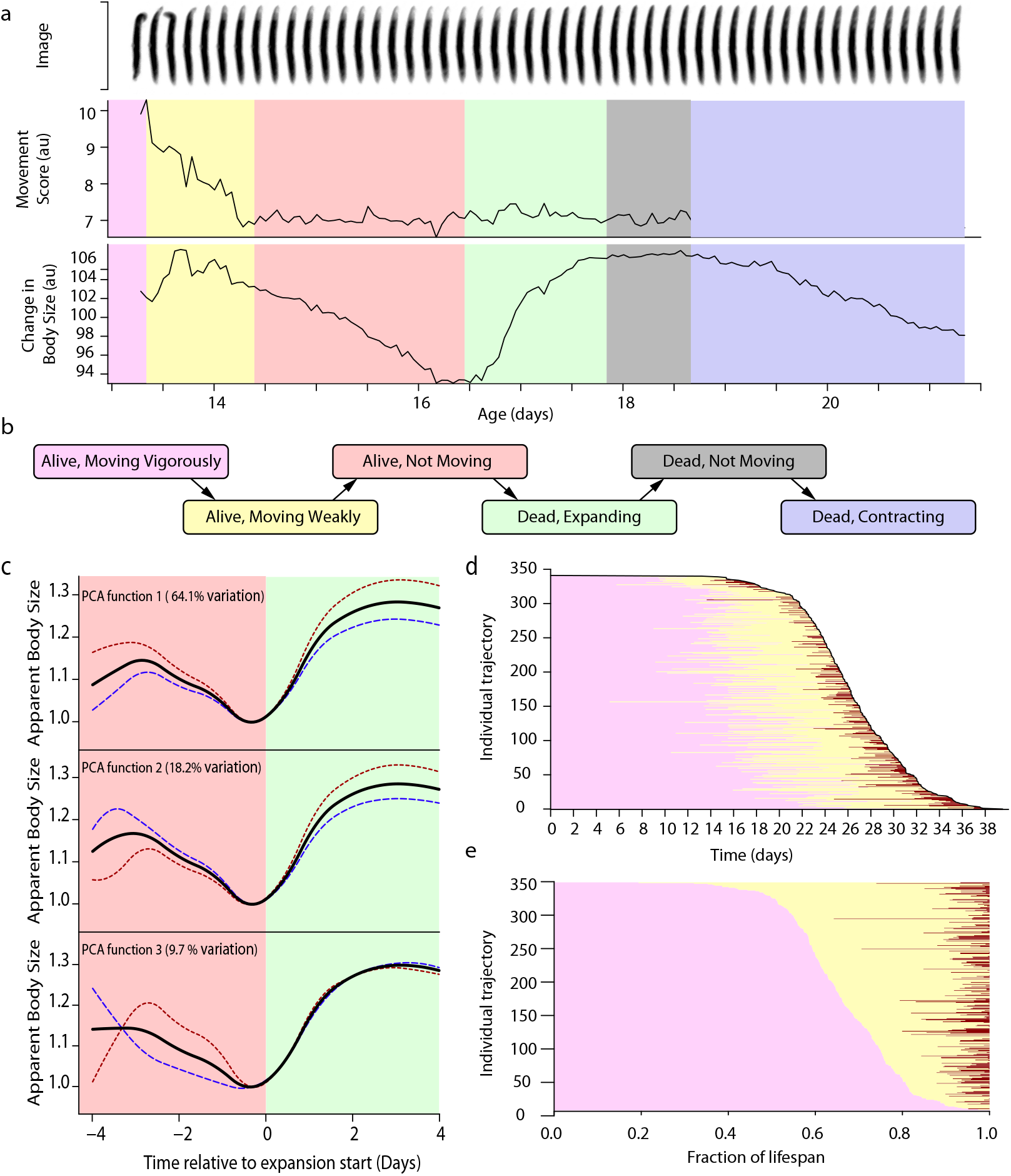
Stages of life and death in C. elegans. **a.** The last few days of a nematode’s life involves a set of stereotypical movement and morphological transitions. After a week or more of moving vigorously *(purple)*, animals start to exhibit only weak postural changes *(yellow)*. After period with no observable movement, during which their apparent body size decreases *(red)*, each individual’s body size increases, plateaus, and then decreases again. **b.** A model of the typical transitions between movement and morphological regimes. **c.** Functional data analysis (FDA) summarizes the variation between individuals in their trajectory of body size near death, as measured in a population of 351 wild-type individuals. The variation was decomposed using principal component analysis (PCA) into three functions, each of which describes an orthogonal aspect of inter-individual variation (+1 standard deviation in red, −1 standard deviation in blue) in relation to the population average trajectory (black). Together, the three functions describe 92% of the total variation within the population. **d.** The time spent in each of the three pre-death states. Each horizontal line represents the state transitions of a single individual. Because the individuals are ordered vertically in descending order, the end of the final state is the death time, and the stacked lines approximate the survival distribution. **e.** The same lifespan data is plotted showing the duration spent in each state as a fraction of each individuals’ lifespan. Individuals are ordered vertically by the fraction of life spent moving vigorously.

To study the variability between individuals in the dynamics and timing of death-associated expansion and contraction, we applied a functional data analytic approach^21^. We fit the trajectory in apparent body size of each individual with a basis spline and performed a principal component analysis of all individuals’ spline parameters (statistical methods). This allowed us to decompose inter-individual variation of trajectories into three orthogonal harmonics^22^ that explain 92% of the variation in the shape of expansion and contraction between individuals. The first harmonic described overall magnitude of both expansion and contraction which explained 64% of the overall variation in shape. The second harmonic described the relative magnitude of contraction and expansion, with some individuals showing smaller contractions and larger expansions, and vice versa, explaining another 18% of variation between individuals. The third harmonic described the rate of contraction prior to expansion, which explained 9.7% of variation. From this, we conclude that though differences exist between individuals in how they contract and expand, these differences have a low-dimensionality. Death-associated contraction and expansion co-occur and despite some internal variability are therefore we can model them as a single event.

### Body-size expansion is a robust marker for nematode death

We then asked whether death-associated contraction and expansion represent a general feature of nematode death, useful as a proxy for death in a broad set of contexts. We used the lifespan machine to observe the lives and deaths of populations of *C. briggsae*, *C. tropicalis*, *C. japonica*, *C. brenneri* and *P. pacificus* nematodes, whose divergent developmental trajectories^23^ reflect a common ancestor living at least 100 million years ago^24^, and repeated the functional data analysis approach to characterize death-associated contraction and expansion. We found that, on average, *C. elegans* and *C. briggsae* show nearly indistinguishable trajectories of death-associated contraction and expansion (Fig 2a). To identify quantitative differences between the two species, we considered the eigenvalues associated with the first PCA harmonic, which describes the shared magnitude of contraction and expansion. Despite inter-individual differences within each species, the overall distribution of magnitudes was indistinguishable between *C. elegans* and *C. briggsae*. (Fig 2.b-c). We then considered the carnivorous nematode *P. pacificus,* which also exhibited a similar shape and magnitude of death-associated expansion and contraction(Fig 2. d-f). We then found that *C. tropicalis*, *C. japonica*, *C. brenneri* strains exhibited qualitatively similar death-associated contraction times, whose typical magnitude (Supplementary figure 2) varied in relation to *C. elegans*. From these results, we conclude that death-associated contraction and expansion are a general feature of death that was likely present in the earliest common ancestor of these species.

**Figure 2:**
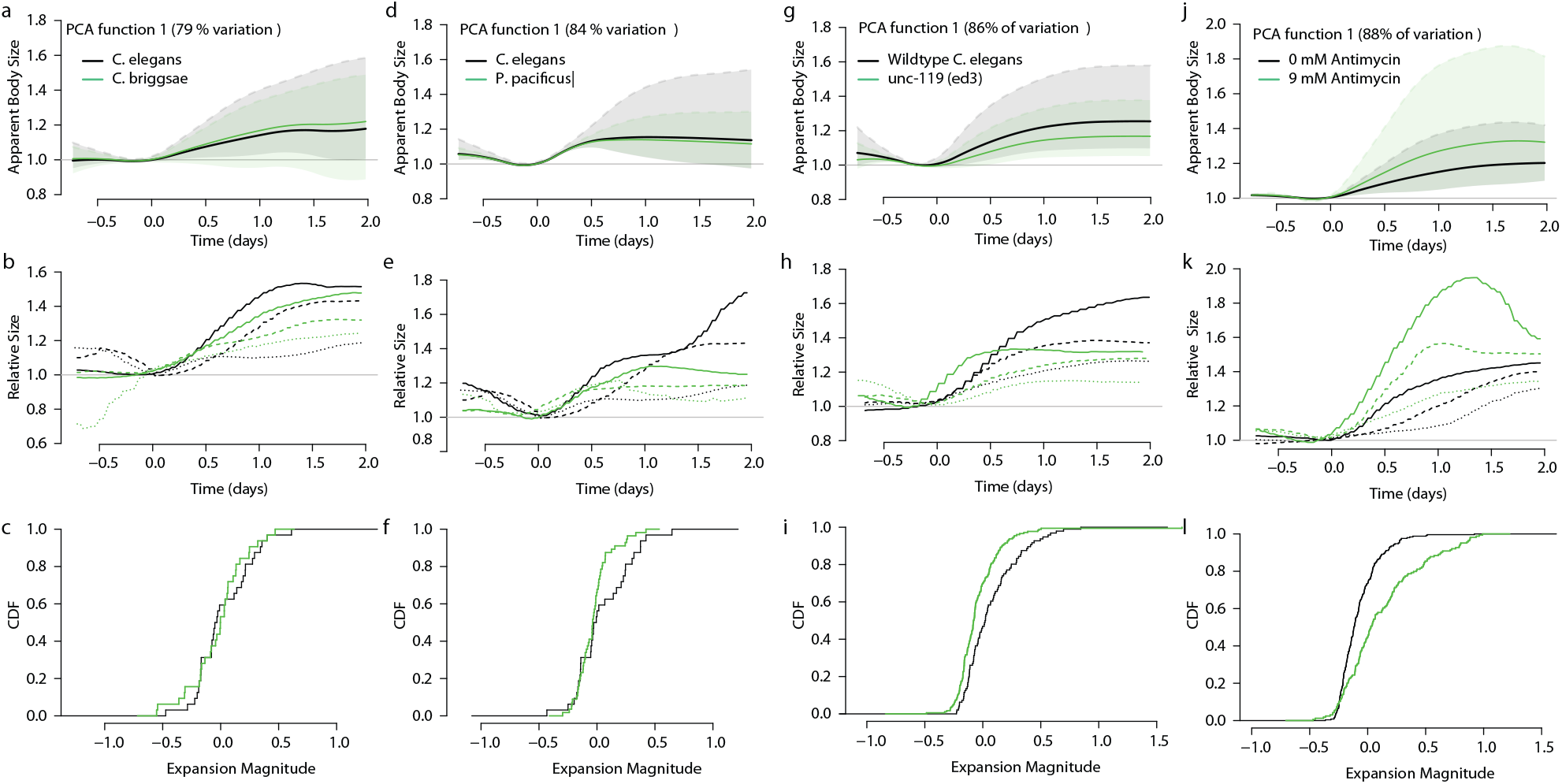
Death-associated morphological changes are conserved across nematode species and paralyzing treatments. **a.** We annotated the movement and morphological transitions by hand for 113 *C. elegans* and 83 *C. briggsae* nematodes. Both strains were analyzed using a functional data analysis (FDA) to identify the largest source of inter-individual variation between trajectories in apparent body size near death, as measured in a population of 351 wild-type individuals. Variation is decomposed using principle component analysis (PCA), and summarized as a single function that depicts inter-individual variation (+1 standard deviation as a dashed line, −1 standard deviation as a translucent line) in relation to the population average trajectory. This allows us to compare both the average trajectory and inter-individual variation between populations of *C. elegans* and *C. briggsae* nematodes. **b.** To provide context for PCA analyses, trajectories are shown for three example *C. elegans (black)* and three *C. briggsae* nematodes of animals, exhibiting the median magnitude death-time expansion *(dashed line)*, a low-magnitude death-time expansion *(dotted line)*, and a high-magnitude death-associated expansion (solid line). **c.** PCA analysis allows us to summarize each individuals’ trajectory as a single “PCA loading” that quantifies the magnitude of an individuals’ trajectory along the PCA function relative to the population average. By plotting the cumulative distribution function (CDF) of PCA scores assigned to individuals of each population PCA allows us to compare the magnitude of death-associated contraction and expansion between *C. elegans* and *C. briggsae*. **d-f** The same PCA analysis was performed to compare 113 C. elegans to 70 *P. pacificus* nematodes, (g-i) 97 wildtype C. elegans to 179 severely paralyzed *C. elegans unc-119 (ed3)* mutants, and (j-l) wildtype *C. elegans* housed either on 0 mM or 9 mM Antimycin A (N=248, N= 166).

To understand the influence of neuromuscular-driven movement on death-associated contraction, we observed the effect of two paralytic interventions: an *unc-119 (ed3)* mutation that interferes with neuronal development and causes severe paralysis^25^, and Antimycin A, an inhibitor of oxidative phosphorylation and severe paralytic agent. Considering a previously published data set^1^, we found that *unc-119 (ed3)* animals showed only a slight decrease in the magnitude of death-associated contraction and expansion (Fig 2. g-i). Treatment with a 9 mM Antimycin A, an inhibitor of oxidative phosphorylation and a strong paralytic, produced an unexpected increase in the magnitude of death-associated expansion (Fig 2. k-l). We conclude that different methods for achieving paralysis can produce different quantitative effects on death-associated contraction, but that even strong paralytic effects do not abolish the phenomenon entirely.

Our data show that death-associated contraction and expansion can provide a, non-behavioral, visible proxy for identifying nematode death. Death-associated contraction is particularly useful for automated methods because, as a non-behavioral event, it does not require external stimulation such as vibration or exposure to blue light^13^ to induce movement.

### Automated identification of movement and behavioral states using HMM models

We then set out to develop automated means for classifying animals’ morphological/behavioral states, and thereby eliminate the laborious by-hand annotation of images. We augmented the lifespan machine software to estimate, in each image collected of each individual, two features describing movement and morphology: an improved “movement score” and a “change in apparent body size” score. The movement score quantifies the magnitude of an individuals’ postural change, calculated as the sum of the absolute value of per-pixel changes in consecutive images of that individual. Apparent body size is estimated as the sum of intensities of all pixels corresponding to an individual in a single image. Because the lifespan machine allows nematodes to freely explore their environment, we found that local features on the agar surface adversely affected our ability to accurately estimate the absolute size of an individual. For example, individuals located in shadows cast by the plate appeared smaller, and animals located on top of inhomogeneities in the bacterial lawn appeared larger. Fortunately, we found that the contribution of such environmental features was time-invariant at the timescale of nematode aging, which allowed us to improve our analysis by focusing on changes in individual body size, which differentiated away errors in absolute size estimation. Therefore, we focused on “movement score” and “change in apparent body size” as features on which to classify animals’ morphological and behavioral states.

To characterize the discriminatory power of our “movement score” and “change in apparent body size” metrics, we compared the distribution of each metric across morphological/behavioral states (Fig. 3 a) in our by-hand annotated wild-type data set. We found that each state exhibited a distinct, characteristic set of distributions of movement scores and change in apparent body sizes (Supplemental Fig. 3 a-b). This suggested that it might be possible to implement a Hidden Markov Model (HMM) to identify morphological/behavioral state transitions from time series of movement scores and change in apparent body size measurements.

**Figure 3:**
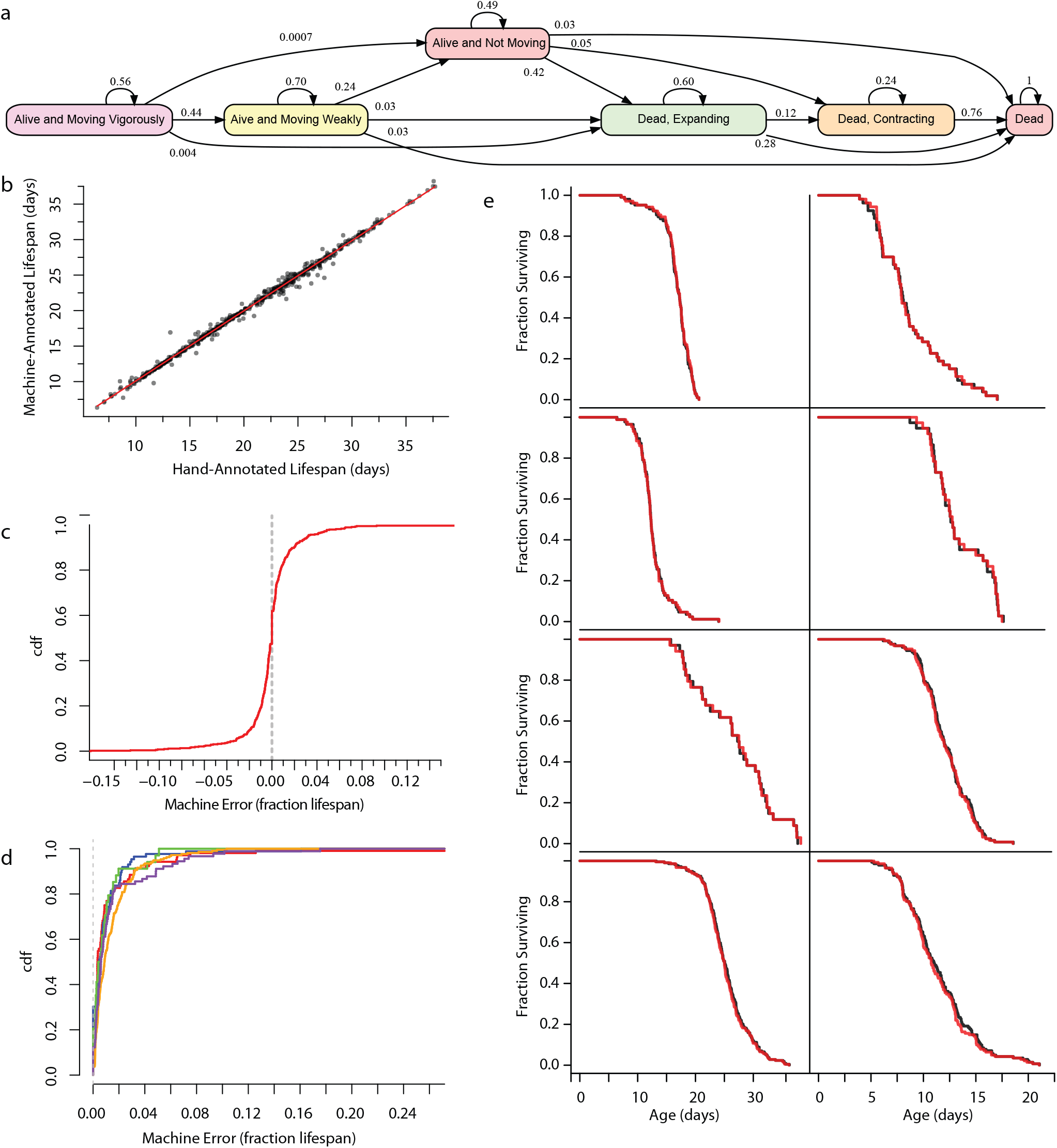
Death-associated morphological changes support robust automated identification of nematode death times. **a.** Using hand-annotated trajectories for 830 wild-type animals whose lifespan were recorded in a variety of environmental conditions, we were able to build a Hidden Markov Chain Model that describes the probability of all possible movement and morphological transitions. The probabilities shown record the probability of an individual making the labeled transition after six hours. **b.** The by-hand annotated data set was sub-divided according to biological replicate and tested in a 6-fold cross-validation approach across biological replicates (statistical methods). Shown is a comparison between the actual and predicted lifespan across all animals in this cross-validation **c.** The cumulative distribution function of machine errors, calculated as the fraction of by-hand lifespan. The same machine errors, but plotted as absolute values and grouped according to the independent biological replicate. **d.** Survival curves comparing the performance of HMM to by-hand annotated data in each of the eight independent biological replicates.

To obtain probabilistic, predictive estimates of movement state, we fit each state’s distributions of “movement scores” and “change in apparent body size” with a two-dimensional Gaussian Mixture Model. To obtain a predictive model of state transition times, we fit the distribution of times spent in each distribution (Fig 1.d) with an exponential model. We then integrated these GMMs using the Viterbi algorithm, which allowed us to models and “solve” an individual’s aging trajectory—identifying the most series of state transition times that best explain the observed time series of movement and body size scores.

To measure the performance of our HMM, we considered a set of 626 hand-annotated, wild-type nematodes chosen from a set of five experiments in which *C. elegans* nematodes were housed in a variety of environmental conditions, including on OP50 lawns, UV-inactivated OP50 lawns, and HT115 lawns and at 20 °C or 25 °C temperatures. A diverse set was chosen to improve the robustness of the resulting models. We used a five-fold cross-validation scheme in which an HMM model was trained on data from four experiments and then used to estimate the state transition times for the remaining fifth experiment. We then repeated this process five times each time using a different replicate for testing. We find that our HMM model estimates show a linear response across a five-week distribution of lifespan (Fig. 4 b) with an average error of 10.9±0.7 hours or 1.4% of lifespan. 95% of individuals had their lifespan estimated within one day of their true lifespan, which corresponds to 95% of individuals being estimated within 4.8% of their true lifespan (Fig. 4 c). Though the HMM model performed slightly better on some experiments compared to others (Fig. 4 d), in no experiment did these errors produce a statistically identifiable difference between the Kaplan-Meier estimates of the survival curves comparing by-hand and automated death times (Fig. 4 e).

**Figure 4:**
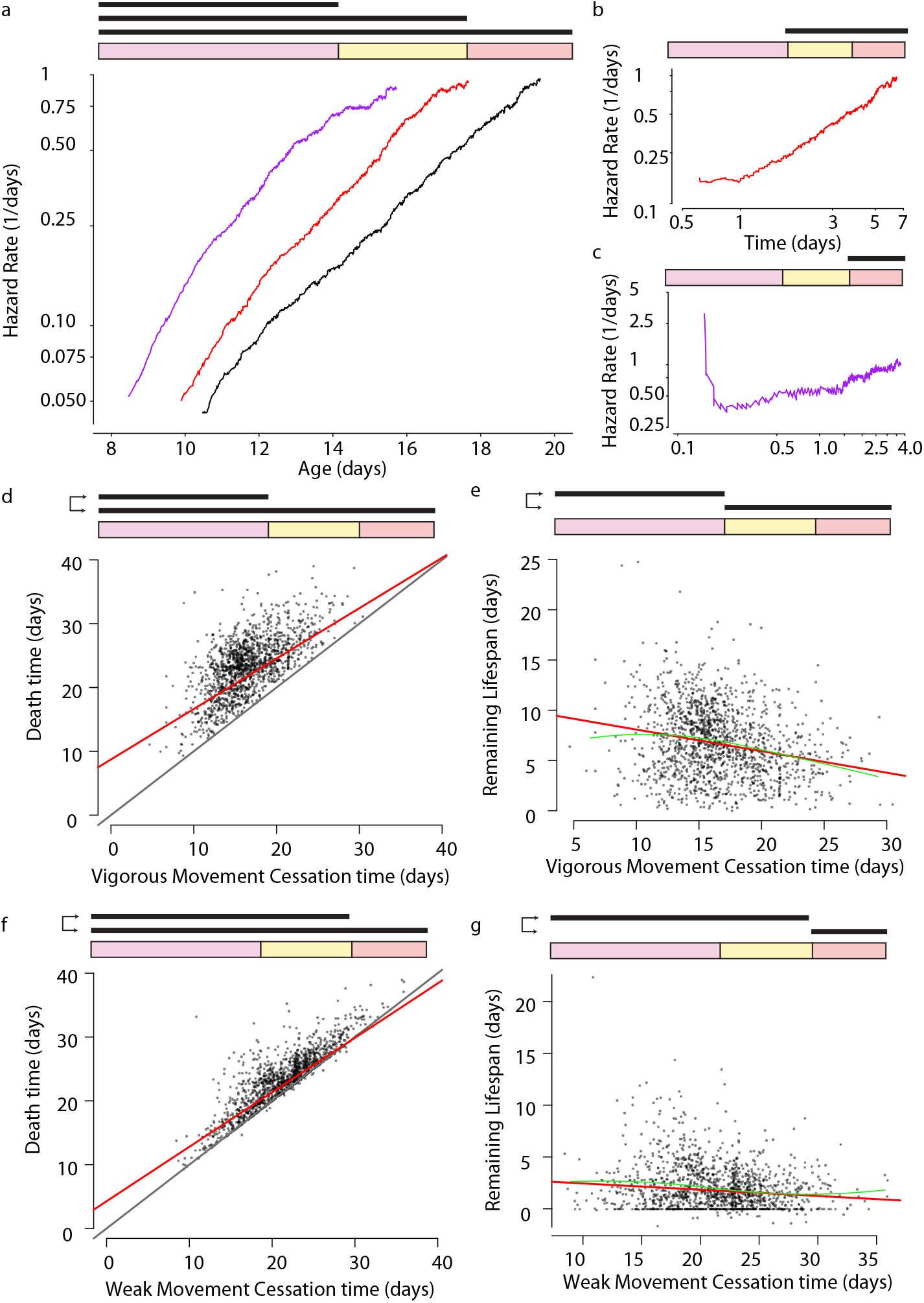
Relating behavioral stages and lifespan in wild-type C. elegans. **a.** In a population of 2346 wild-type C. elegans housed at 25 °C, we compared the risk of animals ceasing vigorous movement (purple), the risk of ceasing all movement *(red)*, and the risk of death *(black)*, all in respect to chronological age. In each panel in this figure, the colored diagrams indicate which behavioral stages are being compared, using the color scheme from figure 1. **b.** We estimated the “clock-forward” risk of an individual’s death relative to cessation of vigorous movement. **c.** We estimated the “clock-forward” risk of an individual’s death relative to the time at which it ceased all movement. **d.** To evaluate the correlation between vigorous movement and death, we compared the timing of vigorous movement cessation to lifespan. The regression line *(red)* (statistical methods) is compared to the unit line *y* = *x (black)*. **e.** Vigorous cessation compared to each individual’s remaining lifespan after ceasing vigorous movement. The regression line *(red)* (statistical methods) is compared to the LOESS regression line *(green)*. **f.** The same analysis as in d, but comparing cessation of weak movement to lifespan and **e.** remaining lifespan.

### The risk of ceasing vigorous movement, ceasing movement cessation, and death rise a polynomial of time

Our new technology allowed us to perform a large-scale, systematic comparison of vigorous movement, weak movement, and death times as they vary across individuals and between populations. We focused first on a population of 2346 wild-type individuals housed at 25 °C, calculating the transition-specific hazard rates that describe the time-dependent risk of an individual transitioning from vigorous movement to weak movement, to no movement and then death. We found that the hazard curves describing cessation of vigorous movement and cessation of weak movement were remarkably similar to those previously observed for death^26–29^, involving a rapid increase in rate starting midlife followed by a gradual deceleration later in life. (Fig. 5a).

**Figure 5:**
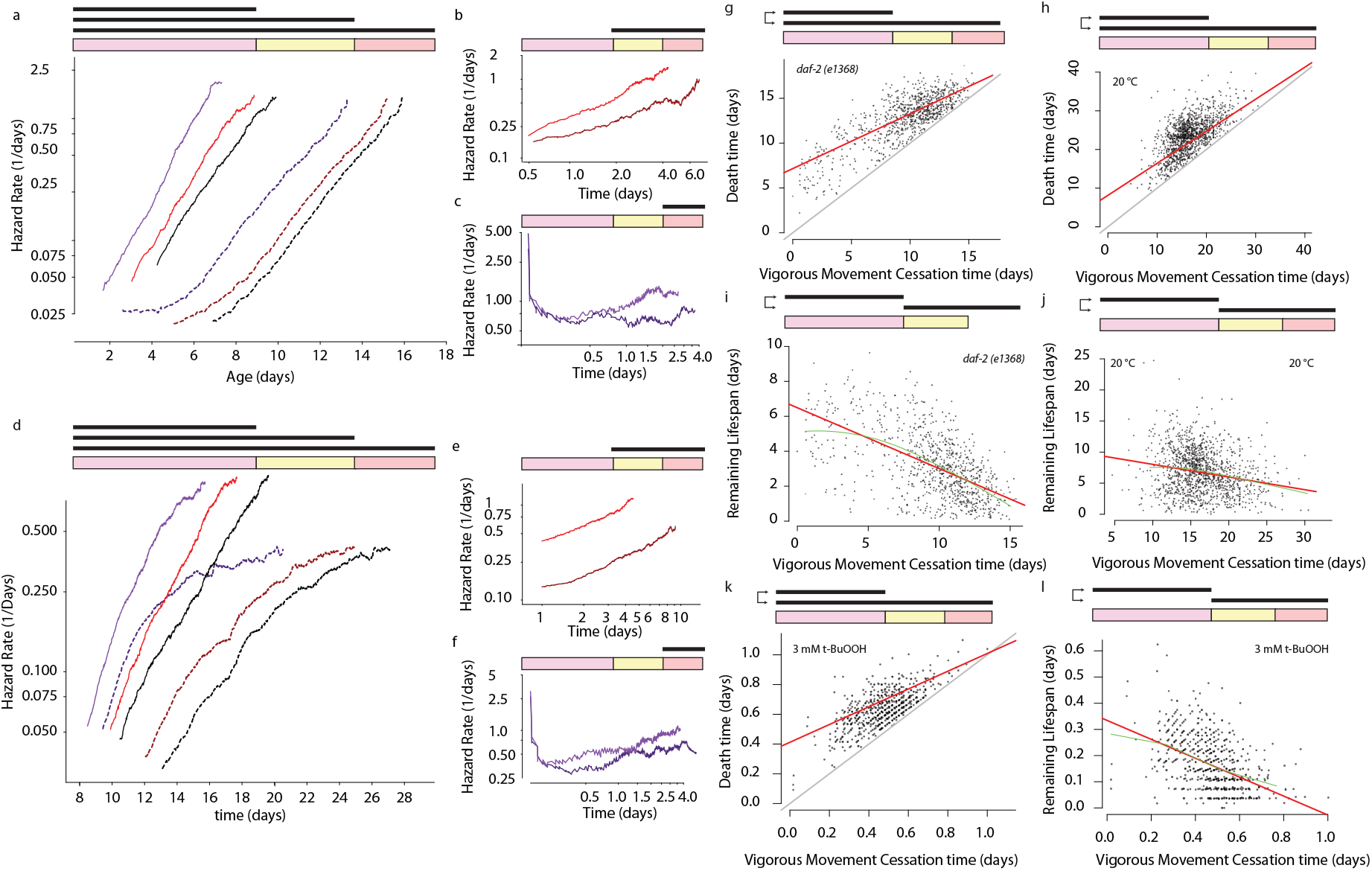
The effect of interventions on behavioral stages and lifespan. **a.** In a population of 919 wild-type and 906 *daf-2(e1368)* individuals, we compared the risk of ceasing vigorous movement (solid purple) ceasing all movement *(solid red)*, and the risk of death *(solid black)* of wild-type populations to the corresponding risks in *daf-2(e1368)* populations *(dashed purple, dashed red, dashed black)*. **b.** We estimated the “clock-reset” risk of an individual’s death relative to cessation of vigorous movement for wild-type populations *(light red)* and *daf-2(e1368)* populations *(dark red)*. **c.** We estimated the “clock-reset” risk of an individual’s death in respect to the time at which it ceased all movement for wild-type populations *(light purple)* and *daf-2(e1368)* populations *(dark purple)*. **d-f.** The same analysis was performed as in a-c, but this time comparing 1441 wild-type individuals housed at 20 °C *(dashed lines, dark colors)* to 2346 individuals housed at 25 °C (solid lines, light colors). **g.**To evaluate the correlation between vigorous movement and death, we compared the timing of *daf-2(e1368)* vigorous movement cessation to lifespan. The regression line *(red)* (statistical methods) is shown in red, compared to the unit line *y* = *x* in black. **h.** We performed the same analysis comparing cessation of weak movement to lifespan. **i.** In a *daf-2(e1368)* population, we compared the timing of vigorous movement cessation to each individual’s remaining lifespan.**i.** The same analysis considering a wild-type population housed at 20 °C **k-l.** We compared the timing of vigorous movement cessation to lifespan and remaining lifespan for a wild-type population exposed to 3 mM t-BuOOH.

To robustly characterize the relationship among vigorous movement, weak movement, and death, we compiled a set of ten independent biological replicates, taken from new or previously published^12,29^ data sets, involving wild-type populations housed on either live bacteria or UV-inactivated bacteria and housed at either 20 °C or 25 °C. These replicates contained on average 810 individuals each. To compare the shape of transition-specific hazard curves across this data set, we explored a set of four parametric distributions: the Gompertz distribution^30^, the inverse Gaussian distribution^31^, the standard two-parameter Weibull distribution^32^ and a three-parameter Weibull distribution with correction for frailty effects^33^. According to the Akaike information criterion (statistical methods), we found that cessation of vigorous movement, weak movement, and lifespan were all best modeled by the three-parameter, frailty-corrected Weibull distribution (Supplementary Fig 4.a-c) a fit previously shown as a best for *C. elegans* lifespan^29^. In three out of ten replicates, the frailty term proved negligible, allowing the two-parameter Weibull to perform equally well as the three-parameter Weibull (Supplementary Fig 4.a), which we interpret as suggesting that our transition-specific hazard functions can be understood as essentially Weibullian with varying amounts of inter-individual variation introduced by population frailty^34^. Compared to death, vigorous movement cessation curves showed lower Weibull beta parameters, reflecting the fact that cessation of vigorous movement always precedes death (Supp. Fig. 4d). Vigorous movement curves also tended to show slightly lower Weibull shape (alpha) parameters (Supp. Fig 4. e) as well as slightly higher frailty (sigma) parameters (Supp. Fig 4f), reflective of a slightly lower slope and more substantial late-life risk deceleration compared to lifespan. The differences in Weibull shape and frailty parameters were sufficient to create statistically significant deviations from temporal scaling in only three of ten replicates. We therefore conclude that vigorous movement cessation, weak movement cessation, and death show qualitatively similar hazard trajectories well described as temporal scaling with deviations of small magnitude only in some replicates.

We then considered the possibility that the risk of an individual’s death might depend not only on chronological time, but also on the time since ceasing vigorous movement. Calculating the “clock-reset”, transition-specific hazard rates^33,35^ (statistical methods), we found that after roughly one day after ceasing vigorous movement, individuals experienced a steady increase in their risk of death (Fig 4.b), suggesting that animals age after ceasing vigorous movement. The risk of death in this period was best described as a two-parameter Weibull distribution (Supp. Fig. 4a-c) with no need for frailty correction. Across all replicates, populations exhibited little or no late-life mortality deceleration in their “clock-reset” hazard curves compared to the chronological age hazard curves (Supplementary Fig. 4 f), reflecting a more homogenous underlying population that aged according to a Weibullian power relationship with time. We interpret this result as demonstrating that a substantial degree of the physiologic variability between individuals responsible for variability in lifespan is established by the time that each individual ceases vigorous movement. As a result, our accounting for vigorous movement cessation times in the “clock-reset” model “aligns” heterogeneous individuals and produces an apparently more homogenous population whose hazard exhibits less frailty-induced deceleration^36^. This observation agrees with longitudinal data^17^ that demonstrated variable individual aging rates early in adulthood to be a major contributor to late-life physiologic heterogeneity. In summary, our “clock-specific” hazard functions show that aging occurs both before and after the cessation of vigorous movement, and that inter-individual heterogeneity reflected in the timing of vigorous movement cessation partially determines the heterogeneity later observed in death times.

We also considered the final behavioral stage in which living animals show no movement. We observed 26% of individuals (618 out of 2346) to die within one hour of ceasing all movement, followed by a four-day period in which the risk of death increased only gradually. It is possible these two phases reflect a biological bimodality present in the underlying aging process. However, this effect can also be attributed to a technical source: at the very end of life, spontaneous movement becomes very slight, perhaps dropping below the measurement threshold of our imaging approach. It may be possible that all animals exhibit some degree of movement right up until the moment of their death that we are only able to observe for three-quarters of individuals. This highlights the importance of using a non-behavioral marker for death for high-precision lifespan experiments, especially in the context of automation. Because this potential technical limitation clouds our interpretation of the non-moving period, in the remainder of this work we focus primarily on vigorous movement and lifespan.

### The duration of vigorous movement is negatively correlated with remaining lifespan

To understand the relationship between vigorous movement, weak movement, and death, we investigated the correlation between the timing of transitions between these stages. Confirming previous observations^18,20,20^, we found that the time at which an individual ceases vigorous movement is strongly correlated the time at which it dies (Fig. 5 d) (partial R^2^ = .45, “statistical methods”). Close inspection of this relationship showed it was well described by a linear model, whose slope we estimated to be significantly less than one (β_v_= 0.91 ; bootstrap p(β_v_ < 1) < 0.0001). This magnitude of slope suggests a systematic phenomenon in which individuals who remain vigorous longer than their peers subsequently spend less time remaining alive, compared to their peers. Indeed, we observed an inverse correlation between the vigorous movement period and remaining lifespan (Fig. 5e) (β_v_= −0.09 ; p (β_v_ != 0) < 0.0001).

Looking for similar effects across our replicates, we observed a linear relationship between vigorous movement and lifespan with a slope less than one (Supplementary Fig. 4 g-i) in seven out of ten replicates. We could not identify an environmental factor that could explain this variability, as all four replicates performed at 20 °C showed slopes significantly less than one, and three out of six replicates performed at 25 °C showed slopes significantly less than one (Supp. Fig. 4k). Exposure to live bacteria also did not explain the variability in slope (Supp. Fig. 4J). So, we then performed a series of analytic calculations and simulations (Supplementary Notes 1-2) to evaluate the performance of our estimation procedure, focusing on the potential impact of environmental variability within experiments and measurement error. We found that environmental inhomogeneity within each experiment—for example, differences in temperature between imaging devices—produced little or no bias in our slope estimates. Furthermore, we found that the magnitude of measurement errors exhibited by our technology and reflected in our data sets was not sufficient to explain the magnitude of slopes we observed. We therefore conclude that some uncontrolled environmental factor, invariant within each replicate but varying between replicates, occasionally increases the apparent slope of the line relating vigorous movement to lifespan.

We then considered the final stage of life, estimating the correlation between the timing of cessation of all movement and death. We observed an even higher correlation (R^2^=.74) between the two events, but no significant deviation from a linear relationship with slope 1 (β_v_ = 0.97; bootstrap p(β_v_ < 1) =0.1), indicating there was no detectible difference in the remaining lifespan of animals who ceased moving early or later in life (Fig. 5g) (β_v_ = −0.03 ; p(β_v_ != 0) = 0.18),

### Interventions alter transition-specific hazard functions but do not alter correlation between vigorous movement and lifespan

To further probe the biological basis of the hazard functions, we considered three interventions known to extend lifespan through a temporal scaling of the lifespan distribution. First, we characterized the disruption of insulin/IGF signaling via the *daf-2(e1368)* allele, known to extend lifespan and extend the duration of youthful vigor^7,8,37^. *daf-2(e1368)* dramatically slowed the age-associated increase in each transition-specific hazard function (Fig. 5a). Using previously published data^29^, we found that the effect of *daf-2(e1368)* on vigorous movement did not reflect a temporal scaling in respect to wild-type animals, as young *daf-2(e1368)* animals were at disproportionately high risk of ceasing vigorous movement, perhaps the result of a subtle quiescence phenotype similar to that observed in stronger *daf-2* alleles^38^. We additionally found that *daf-2(e1368)* individuals experienced a slower rate of aging after vigorous movement cessation (Fig. 5b) as well as a lower risk of death after stopping all movement (Fig. 5c). All of these effects—slower age-associated increases in each hazard function, a slower aging rate after vigorous movement cessation, and a lower risk of death after stopping all movement—were also produced by a second intervention: lowering each individuals’ body temperature from 25 °C to 20 °C (Fig. 5 d-f).

We found that both the *daf-2(e1368)* allele and changes in body temperature did not abolish the linear relationship between vigorous movement cessation time and lifespan. Both populations retained a slope of less than one (Fig. 5 g,h) suggesting that, just as wild-type at 25 °C, the individuals in both populations remained vigorous for longer subsequently died sooner. To exclude the possibility that some time-dependent changes in the environment might increase the risk of death of longest-vigorous moving individuals, we considered a third lifespan-shortening intervention: exposure to tert-butyl hydroperoxide (t-BuOOH). This compound dramatically shortens lifespan, remains stable and potent throughout short lifespan^29^ experiments, and allows us to study animals aging at different rates while keeping other environmental factors constant. We found that t-BuOOH preserved the linear relationship between vigorous movement and lifespan, at both 1.5 mM and 3 mM, with a slope less than one at each concentration(Fig 5.k-l ; Supp. Fig. 5.e) (1.5 mM: β_v_ = 0.85, p(β_v_ < 1) =0.014), 3 mM: β_v_ = 0.60, p(β_v_ < 1) < .0001.

We then expanded our analysis to include five additional interventions—two newly characterized and two characterized using previously published data^12^. We characterized *nuo-6(qm200)* which extends lifespan by disrupting the mitochondrial complex I function^10^, *glp-1(e2141)* which extends lifespan by ablating the gonad*, eat-2(ad1116)*^39^ which extends lifespan by inducing dietary restriction, and changes in diet which in C. elegans can involve a switch from live to UV-inactivated bacteria. We observed a linear relationship between vigorous movement and lifespan on UV-treated and live bacteria, in *nuo-6 (qm200)* populations, and in *unc-119 (ed3)* animals (Supp. Fig. 5g). Animals on Antimycin A exhibited very short vigorous movement periods whose variation we could not measure with accuracy, and so we did not observe or expect to detect an inverse relationship between vigorous movement and remaining lifespan. *glp-1(e2141)* also did not exhibit an inverse relationship between vigorous movement and remaining lifespan, but neither did the wild-type control for that experiment. *eat-2 (ad1116)*, however, appeared to eliminate the inverse relationship between vigorous movement and remaining lifespan, unique in this regard. We consider this result in more depth in a following section.

In summary, we conclude that many lifespan-extending interventions can alter both vigorous movement and lifespan while preserving a negative correlation between vigorous movement and lifespan. This suggests that the negative correlation does not depend on a particular timescale or environmental condition, but instead reflects some structural property of the underlying processes determining vigorous movement and lifespan.

### Environmental factors, diet, and mutations act pleiotropically to disproportionately alter behavioral aging and lifespan

So far, we focused primarily on the nature of interventions’ effect without considering the magnitude of such effects. To compare different interventions’ effect on vigorous movement, weak movement, and lifespan, we applied an accelerated failure time (AFT) regression model. We found that, with the notable exception of paralytic agents Antimycin A and *unc-119(ed3)*, interventions followed a general trend for proportionate effects on vigorous movement, weak movement, and lifespan, evidenced by their alignment along the diagonal in (Fig. 6a-b).

**Figure 6:**
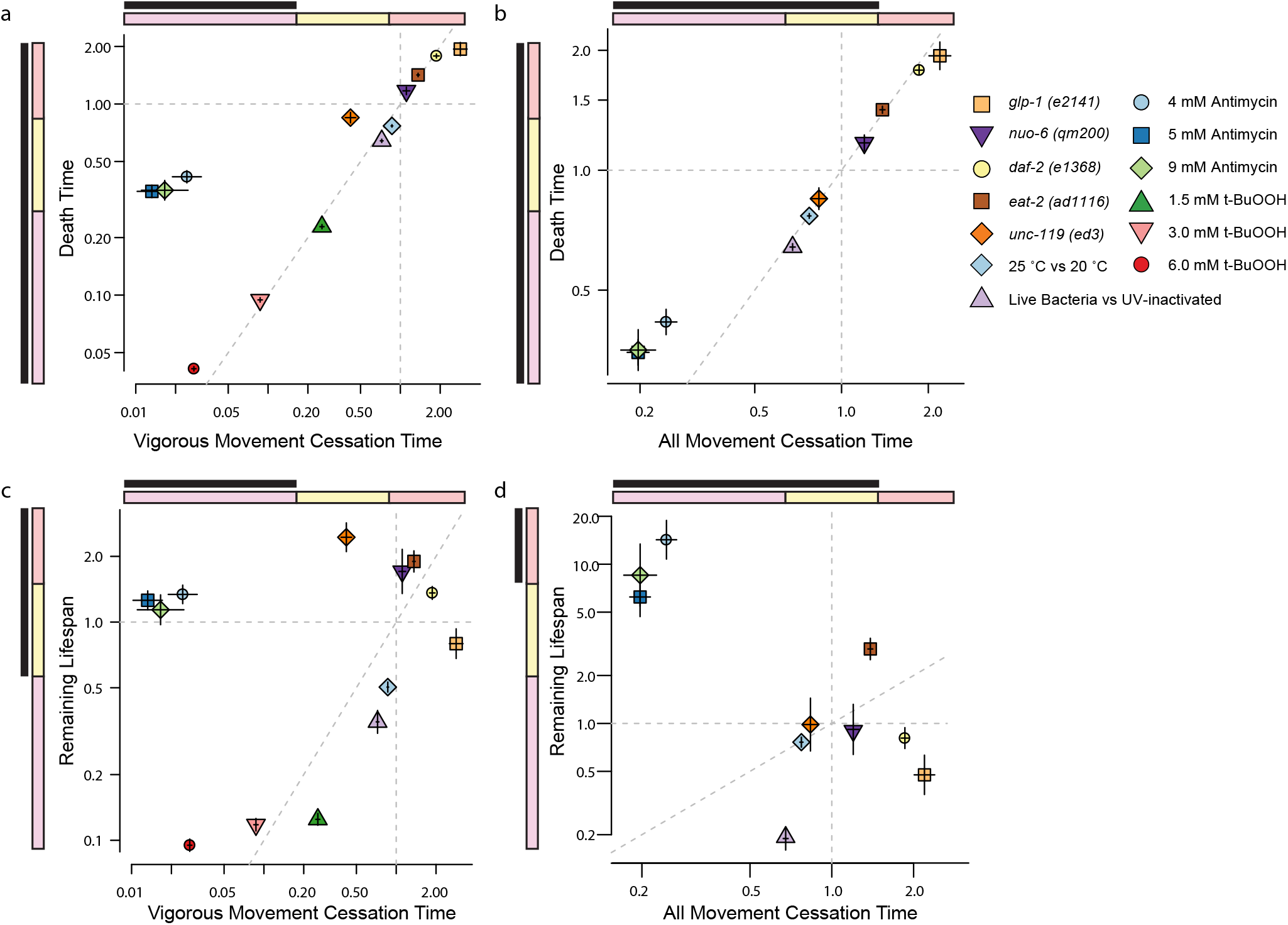
Pleiotropic effects of environmental and genetic determinants of lifespan.. We compared the effect of nine different lifespan-altering interventions on vigorous movement, weak movement, and lifespan. In each panel, the colored diagrams indicate the stages being compared, using the color scheme from Fig. 1. **a.** We compared each interventions’ fold-change effect on vigorous movement cessation and death, relating the AFT regression scale factors observed each event (statistical methods). The interventions tested were *glp1(e2141) (orange square)*, *nuo-6(qm200) (purple inverted triangle)*, *daf-2(e1368) (yellow circle)*, *eat-2(ad1116) (brown square)*, *unc-119(ed3)* (orange diamond), the effect of temperature between 25 °C and 20 °C *(blue diamond)*, the effect of live bacteria vs UV-inactivated bacteria *(light purple triangle)*, 4, 5, and 9 mM Antimycin a *(blue circle)*, *(light blue circle, blue square, green diamond)*, and 1.5 mm, 3mM and 6 mM t-BuOOH. Whiskers indicate bootstrapped 95% confidence intervals. **b.** For the same interventions using the same color scheme, we compared the time at which animals ceased all movement to their death time **c.** For the same interventions using the same color scheme, we compared the time at which individuals ceased vigorous movement to their remaining lifespan, and **d.** we compared the time at which individuals ceased all movement to their remaining lifespan.

This general trend of proportionality involves, however, for each particular intervention a substantial and statistically significant deviation from proportionality (Fig. 6c). The proportionality of the effect of *daf-2* mutation on behavioral aging relative to lifespan has been shown to depend crucially on measurement approach^7,8^ and food source^37^, but here we found that *daf-2 (e1368)* extending both the vigorous movement period and weak-moving period disproportionally more than lifespan (Fig. 8 d,g),. Genetic ablation of the germline via *glp-1(e2141)* had a similar effect, extending lifespan while disproportionately extending the vigorous movement and weakly-moving periods relative to lifespan.

In contrast, dietary restriction induced by the mutation *eat-2 (ad1116)* extended the duration of all behavioral periods while disproportionately increasing the weak-moving and non-moving periods. This result contrasts with a previous study of this *eat-*2 mutant^8^, which found no effect on behavior. We attribute this contrast to differences in the particular behaviors being measured in each study^7^ as well as the increased sensitivity of our method. *nuo-6(qm200)* acted similarly to *eat-2 (ad1116)* in that it disproportionately extended the weak moving period. However, *nuo-6(qm200)* either did not alter or slightly shortened the non-moving period.

Compared to populations at 20 °C, exposure to 25 °C shortened lifespan while having a disproportionately smaller effect on vigorous movement and a proportional effect on the non-moving period. Exposure to live bacteria also shortened lifespan, but produced a disproportionally smaller effect on both vigorous movement and the non-moving periods compared to lifespan. Exposure to t-BuOOH produced a dose-dependent effect, shortening lifespan with a disproportionately smaller effect on vigorous movement at 1.5mM but a disproportionally larger effect at 3 and 6 mM.

*unc-119 (ed3)* shortened but did not eliminate the vigorous-moving period, producing a disproportionate shortening of vigorous movement relative to lifespan while leaving the time spent weakly moving unchanged. Antimycin A similarly disproportionately shortened the period of vigorous movement, but unlike *unc-119(ed3),* Antimycin A dramatically increased the non-moving period relative to lifespan. This suggests that Antimycin A inhibits movement to such a degree that animals cannot detectably move their heads, in contrast to *unc-119 (ed3)* produces a more moderate effect that allows some degree of long-distance movement during early adulthood.

We therefore conclude that interventions frequently produce disproportionate changes in the timing of behavioral state transitions during aging. Pleiotropic effects of lifespan-extending interventions across different aging phenotypes appear to be the norm rather than the exception. However, except for strong paralytics, we saw a general trend in which disproportionate effects arise as small perturbations to a general trend of proportionality, with a course-grained view suggesting that interventions produce approximately proportional effects on behavioral aging and lifespan. We interpret this as suggesting that each interventions’ eccentric, disproportionate effect arises as a quantitative modification to some underlying structure that couples the duration of vigorous movement, weak movement, and lifespan.

### A single intervention—suppression of RNA polymerase II activity—produces different pleiotropies at different doses

To probe the structural relationship between vigorous movement and lifespan, we wished to perform an *in vivo* dosage series involving the quantitative modulation of aging via a single molecular mechanism. Though dosage series have been previously performed by varying environmental factors like temperature, diet, or exposure to tert-butyl hydroperoxide, we hypothesized that an intervention that targeted only a single molecular mechanism might be the least ambiguous approach for relating behavioral aging to lifespan. To realize this goal, we used the auxin-inducible degron system^40^ to obtain long-term, tunable control of aging. Focusing on the RNA Polymerase II complex subunit *rpb-2*, we reasoned that because RPB-2 is required for normal lifespan (Supplementary Fig. 7a), a quantitative degradation of this protein might produce a quantitative shortening of lifespan. Editing the endogenous locus of *rpb-2* to introduce a short degron tag^41^, we obtained a strain that allowed us to selectively ubiquitinate and degrade RPB-2 using the auxin-inducible Arabidopsis TIR1 E3-ubiquitin ligase transgene. We hypothesized that chronic exposure to different auxin dosages would produce differences in the steady-state degradation rate of *RPB-2::AID* and thus yield different average activity levels of the RNA Polymerase II complex. By exposing populations to a dosage series of auxin, we predicted that we could obtain a dose-response curve linking quantitative changes in *RPB-2 activity*, precisely capturing its quantitative influence on both behavioral aging and lifespan.

Testing this production, we observed that the synthetic auxin analog ɑ-Naphthaleneacetic acid (NA)^42^ produced a dose-dependent, monotonic decrease in the lifespan of our *rpb-2::AID* strain (Fig. 7a). At 8 mM NA, a relatively high dosage, we found that *rpb-2::AID* animals lived on average 81% shorter compared to unexposed animals. In contrast, control animals lacking *rpb-2::AID* lived only 16% shorter, indicating a small degron-independent effect of NA on lifespan.

**Figure 7:**
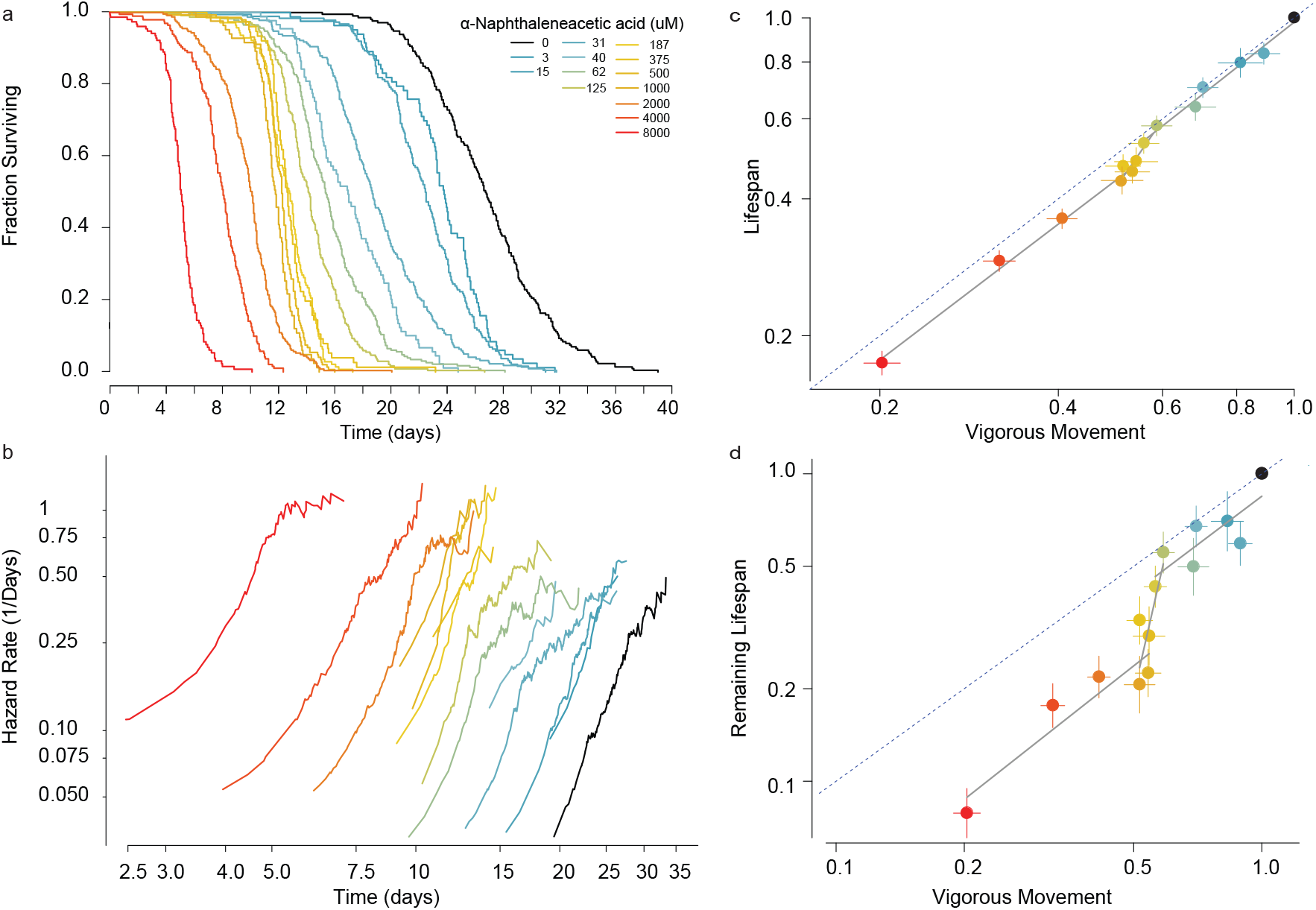
An RNA Polymerase II dosage series. **a.** A transgenic *C. elegans* strain allows tunable degradation of the RNA polymerase II subunit RPB-2 using the auxin-inducible degron system. Kaplan-Meier survival curves demonstrate that increasinge concentrations of alpha-Naphthaleneacetic acid (NAA) shorten *C. elegans* lifespan. **b.** The time-dependent trajectory of the risk of death was calculated for animals exposed to each dosage. **c.** At each dosage, we compared the effect of NAA on the time of vigorous movement cessation and lifespan. Data was fit by a tri-phasic linear model (grey lines; statistical methods). **d.** We performed the same analysis, but this time comparing vigorous movement cessation to remaining lifespan.

We then considered the effect of RPB-2 knockdown on each population’s transition-specific hazard function. We found that NA exposure increased the rate of increase in the risk of animals ceasing vigorous movement as well as the rate of increase in their risk of death (Fig. 7b). We interpret this as suggesting that increased RPB-2::AID degradation influences vigorous movement and lifespan primarily by modulating some underlying rate or rates of aging.

We then considered the effects of RPB-2::AID on the time that individuals spend moving vigorously relative to their lifespan. We observed a triphasic RPB-2::AID dose-response curve (Fig. 7c) that was well-represented as a piecewise linear effect. Compared to wild-type animals, we found that RPB-2 degradation produces a disproportionately greater effect on vigorous movement relative to lifespan. However, at all concentrations below 125 μM, we observed RPB-2 to produce a proportional effect on vigorous movement and lifespan, maintaining a constant ratio between the two even while both are shortened by increasing NA dosages. We identified an interval between 125 μM and 500 μM in which increasing dosages of NA had a negligible effect on the vigorous movement stage, but progressively shortened lifespan. At concentrations higher than 1 mM NA, RBP-2 degradation again produced a proportional effect on vigorous movement and lifespan, but this time maintaining a new but constant ratio.

We interpret this segmental dose-response as evidence for a hierarchical structure linking RPB-2, behavioral aging, and lifespan. At concentrations less than 125 μM and above 500 μM, RBP-2 produces an equal effect on both vigorous movement and lifespan, suggesting it influences both through a common upstream mechanism. In contrast, between 125 μM and 500 μM, RPB-2 produces a differential effect on vigorous movement and lifespan, suggesting it can influence each separately. Above 500 μM, RBP-2 again influenced both equally. We therefore conclude that auxin-induced RPB-2 degradation can effect behavioral and lifespan both through shared and independent mechanisms.

How might this effect arise? It might be that behavioral aging and lifespan have physiological requirement for RBP-2 activity in each dosage interval. For example, in animals with unperturbed RPB-2 activity, normal vigorous movement might require more RPB-2 activity than does lifespan, meaning that small decreases in RPB-2 depletion shortens vigorous movement more than lifespan. Then, as the degree of RPB-2 depletion increases, a severe shortage of RPB-2 might rewire physiology to alter the relative dependence of vigorous movement and lifespan, and thereby change the proportionality of RPB-2’s effect in higher dosage intervals. Another model would explain the segmental dose response curves as arising from some “off-target” action of NA that arises only at high doses. Such an “off-target” effect might nevertheless might be a useful model for understanding the pleiotropic influence of more conventional lifespan-extending interventions, as any intervention might be expected to produce increasingly severe “off-target” effects as its intensity is increased.

In summary, we find that RBP-2 suppression via NA produces pleiotropic effects on behavioral aging and lifespan whose relative magnitude is fixed within dosage intervals, but changes between dosage intervals. We arrive at a similar conclusion as before, when considering multiple different interventions in Fig 6: Lifespan-altering interventions can act disproportionately to alter the period of vigorous movement and lifespan, but such disproportional effects arise as a modulation of some underlying structure that somehow couples vigorous movement and lifespan, leading to a general trend of proportional effects.

### A hierarchical process model for vigorous movement and lifespan

Any model to relate vigorous movement and lifespan must explain four findings: 1) the positive correlation observed between vigorous movement and lifespan, 2) our estimates of the linear model between vigorous movement showing a slope of less than one, implying a negative correlation between vigorous movement and remaining lifespan 3) the action of each intervention to produce eccentric disproportionate effects on vigorous movement and lifespan 4) the general trend across interventions to produce approximately proportional effects on vigorous movement and lifespan.

This first finding, a positive correlation between vigorous movement and lifespan, is widely interpreted as suggesting that behavioral aging shares some upstream determinant in common with lifespan. This determinant appears to be shared among many additional life history traits including declines in pharyngeal pumping rates and reproductive span^20^. The most straightforward explanation would has vigorous movement cessation and lifespan arising as distinct manifestations of a single aging process. In this way, vigorous movement would report on an individual’s “health” or “biological age” that measures the distance that individual has traveled along some path from youth to death. However, this model cannot alone parsimoniously account for observed linear relationship between vigorous movement and lifespan with a slope less than one, nor the associated negative correlation we observe between movement span and subsequent lifespan. Were vigorous movement cessation and death two steps of the same process, then some additional phenomenon would be required generate the negative correlation between vigorous movement and remaining lifespan (Supplementary Note 1). For example, it has been shown that in *Drosphilia melanogaster*, increased exercise during youth leads to a shorter lifespan^43^ suggesting some tradeoff between youthful vigor and lifespan. However, this negative relationship between populations does not necessarily involve a negative correlation between individuals within a population, and furthermore the effect of exercise appears to be beneficial and lifespan-extending many other species including, rodents^44^, humans^45^, and *C. elegans*^46^. Therefore, lacking evidence fora physiologic mechanism that would introduce a negative correlation between vigorous movement and lifespan, we set out to identify a model that would innately produce such a correlation.

A negative correlation between vigorous movement and remaining lifespan would arise naturally if we discard the presumption of a causal relationship between the two stages. If we instead allow the positive correlation between the two stages to arise from the confounding influence of a shared upstream factor. This upstream factor would need to vary between individuals and have a similar effect on both: extending the vigorous span and lifespan to the same degree. This would allow the magnitude of the slope between the two spans to take a value between zero and one, depending on the magnitude of influence from the shared upstream factor. A slope of one would suggest the upstream factor fully determines the timing of both spans, and a slope of zero would imply that no upstream factor has any influence on both spans. Therefore, in a hierarchical process model, the negative correlation between vigorous movement and remaining lifespan would arise as an epiphenomenon produced by the subtraction of two partially dependent random variables. This argument is derived formally in Supplementary Note 1.

### Simulating the hierarchical process model

To demonstrate the behavior of the hierarchical process model, we performed a stochastic simulation. Adopting the assumptions of previous work^31^, we considered a set of individuals who begin “life” all sharing an equivalent physiologic state, with each individual then experiencing a decline in this state according to a biased random walk (Weiner process). State transitions—between vigorous movement and death—are defined to occur at the time each individual passes below a certain distance threshold, with subsequent thresholds defining subsequent states. This type of Weiner process model, in the single-threshold case, has been shown to produce mortality statistics that recapitulate many aspects of biological aging^31^.

In this Weiner Process simulation, we observed no correlation between state transition time and the duration of time spent in the subsequent state (Fig. 7a), as expected for a memoryless random walk. So, we introduced a frailty term^34,47^ into the model, allowing each individual to have its own characteristic drift rate such that some individuals progressed faster on average than others. This frailty mimics the heterogeneity in aging rates observed in populations of *C. elegans* and humans, where indistinguishable individuals exhibit consistent, life-long differences in the risk of adverse age-associated events and visible aging phenotypes^11,17^. Adding frailty to our model produced a positive correlation between state transition time and subsequent state duration (Fig. 7b), demonstrating the ability of a confounding factor to generate apparent correlations among the multiple passage times of an otherwise memoryless process. Our simulation, however still did not recapitulate the negative correlation we observed in our empiric data, between cessation of vigorous movement and remaining lifespan.

One way to recapitulate a negative correlation between vigorous movement and lifespan involves changing the nature of frailty such that it produced paradoxical effects in different life stages. We considered a paradoxical frailty model, in which animals with an individual aging rate lower than their peers early in life subsequently reversed their population-relative rate to age more quickly than their peers later in life. Such a model, by construction, produced a negative correlation between vigorous movement and remaining lifespan (Fig. 7c).

However, our analytic calculations suggested a negative correlation could arise parsimoniously, without some mechanism for generating paradoxical frailty effects, if we discarded the assumption that vigorous movement and lifespan arise as sequential steps of some underlying process. Instead, we allowed the two stages to be determined by two independent Weiner processes. The resulting model then produced uncorrelated state transition times, but also generated a negative correlation between vigorous movement and lifespan (Fig. 7d). This negative correlation raises from a basic property of independeny random variables, such that the quantity Y-X is always inversely correlated with X with a slope of −1. This model however, still failed to recapitulate our experimental data, as it failed to produce a correlation between vigorous movement and lifespan.

Finally, we added a frailty effect the two-process model. The model then recapitulated the positive correlation between vigorous movement and lifespan with a slope of less than one, as well as the negative correlation between vigorous movement and remaining lifespan with a slope less than zero (Fig. 7e). In conclusion, our simulations validate our analytic characterization of the hierarchical aging model (Supp. Note 1), that showed the model should reproduce the statistical relationships we observe in our data.

### Multiple aging processes need be only partially independent

We performed one final simulation, to further characterize the hierarchical process model. We modified the two-process frailty model (Fig. 7e) to allow a tunable dependence between processes (Sup. Fig. 7). As before, at the start of each simulation random walkers start at the same position and then decline following to two biased random walks A and B, with walk A (blue) determining the transition time into state 2 (weak movement), and walk B (red) determining the transition time into state 3 (death). However, at each step of the simulation the two random walkers progress according to the weighted sum of a shared random variable and a process-specific random variable. By altering the weight of this weighted sum, we were able to tune the exact degree of independence between processes A and B. We then measured the effect of this dependence on the correlation structure of transition times between states. As predicted, the linear relationship between transition times took a value between 0 and 1 that increased with the influence of the shared upstream factor (Supplementary Figure 6). We conclude, in the hierarchical process model, the slope of the linear relationship between vigorous movement and lifespan reflects the degree of independence between the processes determining vigorous movement and lifespan. Future efforts to obtain empiric estimates of all parameters of in the hierarchical model may provide a means to estimate the degree of independence between biological aging processes, allowing us to predict the effect of pleiotropic interventions in ageing.

## Discussion

In this work, we improved the Lifespan Machine and applied it to study the pleiotropic action of lifespan-extending intervention. This technical development, involving the characterization of a evolutionary-conserved, non-behavioral marker for death, allowed us to systematically investigate the relationship between behavioral aging and lifespan. Dividing lifespan into vigorous moving, weak moving, and non-moving periods, we confirmed the previously identified positive correlation between vigorous movement cessation and lifespan^6,7,17,18,20^ and then discovered that the slope of the linear model relating these two events takes a value less than one, reflecting a negative correlation between vigorous movement duration and each individual’s remaining lifespan.

Using analytic calculation and simulation, we demonstrated how such correlations need not reflect a causal relationship between vigorous movement and lifespan, nor a tradeoff between the aging rate early and late in life. Instead, the observed correlations arise parsimoniously from the two aging processes (Fig. 10) under the influence of a shared upstream factor. This hierarchical model allows each individual to have two distinct “biological ages”, one describing a decline in the physiology required to move vigorously and the other describing a decline in the physiology needed to remain alive. With two physiologic processes declining in parallel under the influence of some shared upstream factor, a linear relationship between them would be expected to take a value less than one, and a negative correlation between vigorous movement and remaining lifespan would arise as an epiphenomenon.

A hierarchical model would also explain the multiscale results of our interventional experiments. Across diverse interventions and RPB-2 dosage series, we identified a coarse-grained tendency towards proportional effects on vigorous movement and lifespan. Yet, inspecting each intervention individually, we observed substantial deviations from proportional effects. We believe that this multiscale behavior is the natural result of interventions having two actions: a direct influence on the two processes determining vigorous movement cessation and lifespan combined with an indirect influence some shared upstream factor. Then coarse-grained proportionality would from the action of interventions on the shared upstream factor, with individual disproportionalities arising from the direct effect of interventions on each process. Such an organization is reminiscent of the classical clinical definition of frailty^48^

A hierarchical aging model also matches the organizational structure of multicellular aging. *C. elegans*, mice, and humans similarly experience aging as a complex interplay between physically distinct but systemically coupled cells, tissues and organs. The separation of physiology into distinct compartments—cells, tissues, organs—provides an obvious mechanism by which multiple aging processes could proceed under shared systemic influence. While tissues remain physically separated, they nevertheless co-exist within a single individual, and therefore share a host of hormonal, structural, and environmental influences that would provide an obvious mechanism for coupling aging rates across tissues. We speculate that if muscle, intestine, and nerves were to decline in distinct ways over time, with vigorous movement depending mostly on muscle and nerves but lifespan depending mostly on nerves and the intestine, then this overlapping organ-level dependence could easily result in the emergence of multiple, partially dependent aging processes. Just as organs represent spatially distinct physiologic compartments under the influence of shared systemic factors, aging might represent the outcome of distinct dynamical processes under the influence of shared systemic factors.

Our data and framework also provides additional context for the concept of “healthspan”, which in *C. elegans* humans is often defined using youthful vigor as proxy for biological age. Our findings highlight the pitfalls of using behavioral measurements as a proxy for some single organismal state called “health” assumed to be somehow causally involved in determining lifespan. We find that decrepitude and death are not a consequence of reaching the end of one’s behavioral healthspan. Instead, our findings suggest that good behavioral health and longevity are experienced not sequentially but in parallel as distinct conditions driven by partially dependent underlying physiologic declines. It could be a mistake to assume that youthful vigor reflects health in any fundamentally predictive sense, as we have shown that interventions often decouple youthful vigor from lifespan. Instead, our model suggests that individuals posess multiple biological ages, each describing an animal’s location along an independent process of physiologic decline, that exhibit complex interdependencies. And if individuals have multiple biological ages, then they must by definition have multiple “healths” as well.

Our model suggests that the interpretation of biomarkers to infer biological age must be done with great care, recognizing the potential complexities of partial dependencies among multiple aging processes. In a complex hierarchical system, one risks a linguistic quagmire when using the term “health” to generalize from any particular aging phenotype. It will be fascinating to see the results of clinical trials of anti-aging therapies currently underway. Our model would predict a rich in the consequences of clinical intervention, involving at least two potentially separable effects: a direct effect of interventions on specific aging processes combined with an indirect affect mediated by shared upstream factors. In particular, recent advances in measuring tissue-specific histone methylation markers^49^ may provide a means for identifying and separating the direct and indirect effects of clinical interventions in aging.

## Author Contributions

NO developed auxin-inducible *rpb-2* degron strains, optimized ɑ-Naphthaleneacetic preparation and dosage strategies, and performed experiments. SS designed and calibrated imaging equipment, and performed experiments. MAMB performed experiments and constructed and calibrated equipment. OMFM implemented, tested, and optimized image acquisition software. NS conceived and designed experiments, designed and implemented software, performed experiments, analyzed data, interpreted results, and wrote the manuscript.

## Materials and Experimental Methods

All animals were housed at 20 °C or 25 °C at a density of approximately 40 animals per agar plate, and either live OP50 bacteria or UV-inactivated NEC937 (OP50 ∆uvrA; KanR) ^50^. Fig. 1,5,6, and 9: 20 °C UV-inactivated; Fig. 8a: 20 °C; Fig 8b: UV-inactivated; Fig 8c,d: 25 ° UV-inactivated; Fig. 2 and 8e,f 25 ° Live. F In all experiments, age synchronous populations were obtained by hypochlorite treatment of gravid populations. To eliminate progeny, populations were transferred at late L4 stage onto plates containing 27.5 μg ml^−1^ 5-fluoro-2-deoxyuridine (FUDR, Sigma) to eliminate live progeny. Where live bacteria were used, 10 μg ml^−1^ FUDR sufficed to eliminate live progeny. 1-Naphthaleneacetic acid (Sigma-Aldrich), was solubilized in 1M potassium hydroxide and then added into molten agar. *rpb-2* RNAi in Supp. Fig 9 was performed using age-synchronous spe-9(hc88) I; rrf-3(b26) animals housed at 25 °C in the absence of FUDR, transferring animals at L4 stage from empty vector HT115 onto HT115 bacteria containing the *rpb-2* construct.

**Figure 8:**
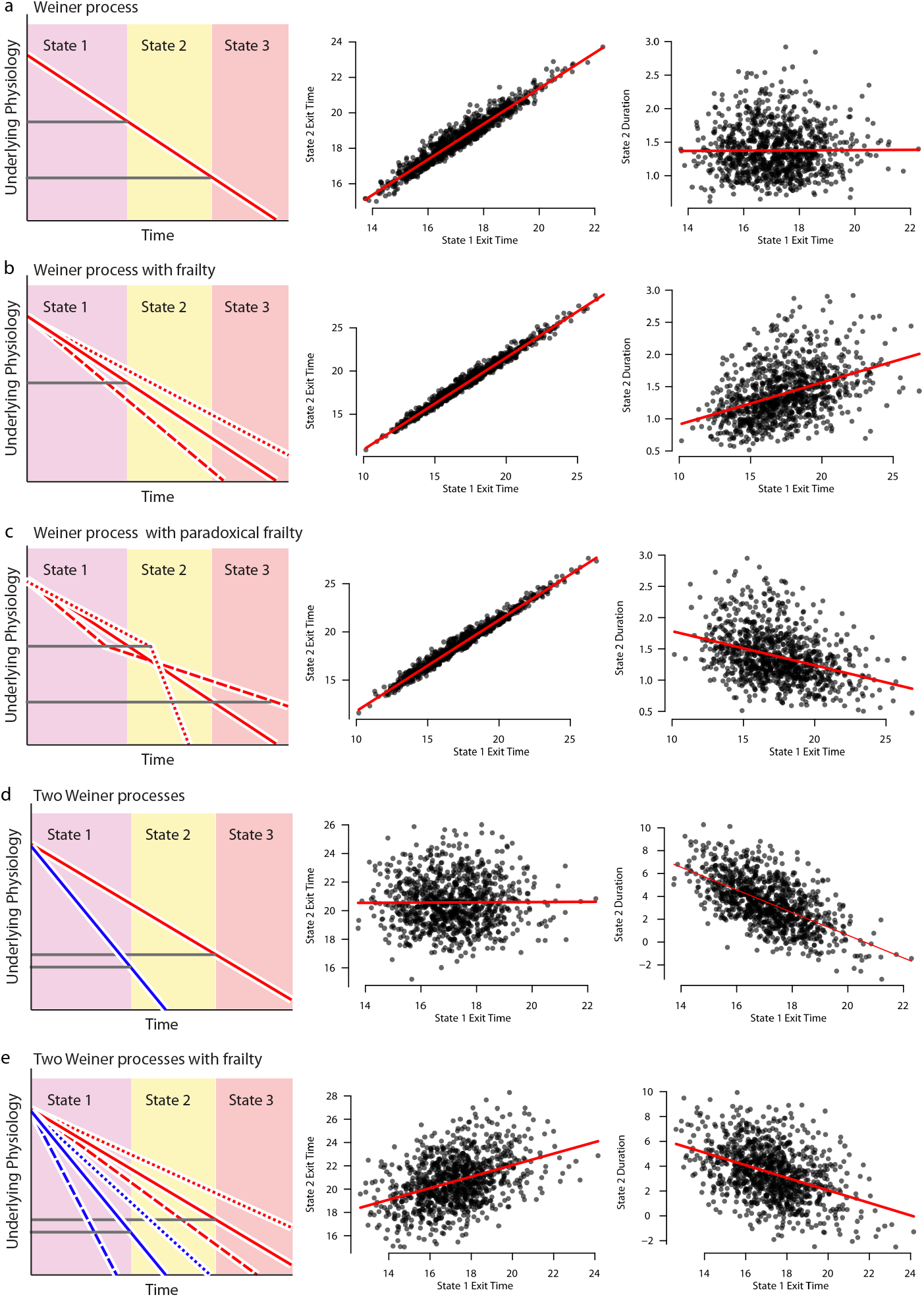
Stochastic models link physical declines to behavioral state transition times. To understand our empiric behavioral state transition data, we considered a series of stochastic simulations modeling biased random walks (Weiner processes). In each panel, a diagram *(left)* shows the specific model being simulated. For each simulation, the relationship between state transition times is plotted *(middle)* and fit via linear regression *(red)*. Then, the relationship between the first state transition and the second state duration is plotted *(right)* and fit via linear regression *(red)*. **a.** A Weiner process, in which each state transition is determined by the sequential crossing of two thresholds. **b.** A single Weiner process in which a frailty effect causes individuals to vary in their drift rate. **c.** A single Weiner process model with a paradoxical frailty effect, where individuals vary in their drift rate to equal and opposite degrees before and after entering state 2. **d.** A two process Weiner model, in which the exit from state 1 is determined by one process *(red)* and the exit from state 2 determined by the other (blue). **e.** A two-process Weiner model including a non-paradoxical frailty effect.

**Figure 9:**
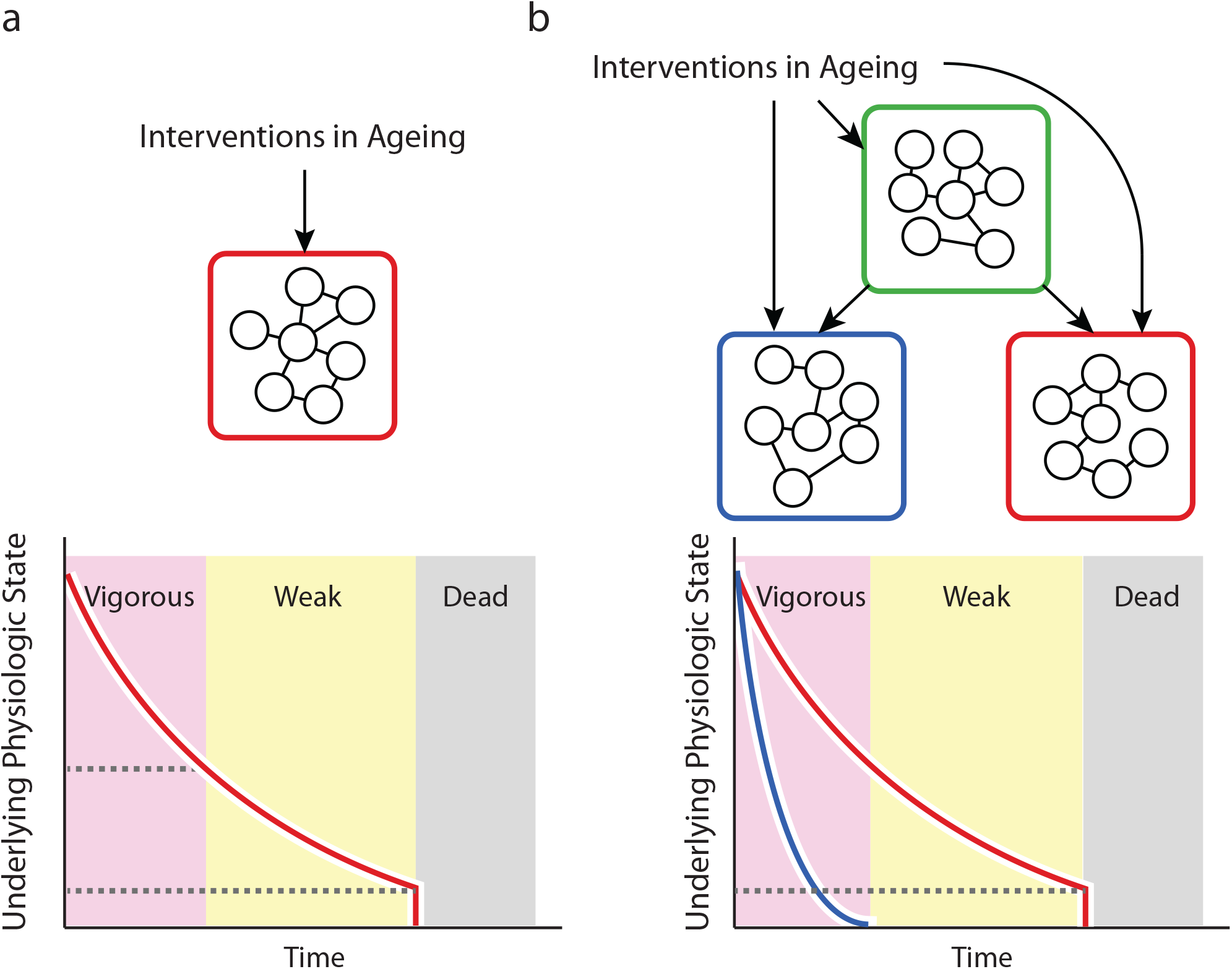
A hierarchical model of vigorous movement and lifespan. **a.** Our data are not parsimoniously explained by a model in which behavioral stages and death emerge as sequential manifestations of a single underlying aging process. Here, the combined function of all physiologic components in the red box declines over time, as shown by the red line the graph. Cessation of vigorous movement and death are determined when the components’ function declines past two sequential thresholds *(dotted lines)* **b.** Our data suggest a hierarchical process model in which behavioral stages and death are determined by distinct aspects of physiology. Functional declines in the determinants of vigorous movement *(blue)* decline in parallel with the determinants of lifespan *(red)*. Both declines are influenced by a shared upstream factors *(green)*. The direct effect of an intervention on the blue or red boxes can be distinguished from broader, systemic effect mediated by the green box, based on the correlation structure of vigorous movement and death times.

By hand annotation of individual worm trajectories used in Figs. 1–4 were performed using the Worm Browser, which is part of the Lifespan Machine software package available at https://github.com/nstroustrup/lifespan.

## Acknowledgements

We thank J. Alcedo (Wayne State University) for nematode strains, X. Manière (Université Paris Descartes) for providing the NEC937 *Escherichia coli* strain used for UV-irradiation, Javier Apfeld, Tom Kolokotrones, Sebastian Maurer, Elvan Boke, and Ben Lehner for constructive feedback on the manuscript, and all members of the Stroustrup laboratory for discussions and encouragement throughout the project. We thank Yan Qi and Winston Anthony for establishing antimycin A assays on the Lifespan Machine. Some strains were provided by the CGC, which is funded by NIH Office of Research Infrastructure Programs (P40 OD010440). This project has received funding from the European Research Council (ERC) under the European Union’s Horizon 2020 research and innovation programme (Grant agreement No. 852201), the Spanish Ministry of Economy, Industry and Competitiveness (MEIC) to the EMBL partnership, the Centro de Excelencia Severo Ochoa, and the CERCA Programme/Generalitat de Catalunya. This work was supported by the MEIC Excelencia award BFU2017-88615-P, and by an award from the Glenn Foundation for Medical Research.

---

**Supplementary Figure 1:**
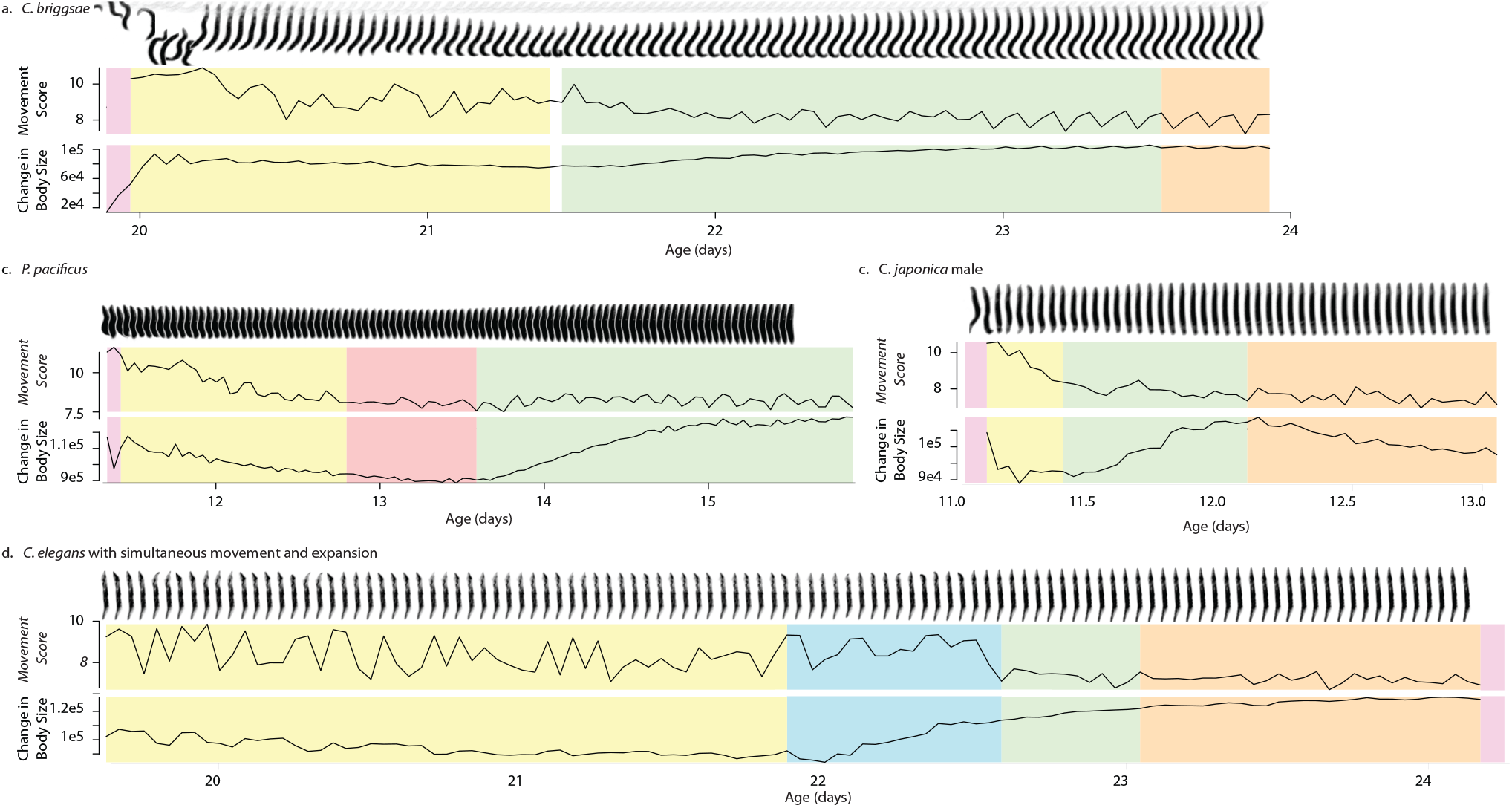
Stages of life and death in *C. elegans.* **a.** The death-associated contraction and expansion of a *C. briggsae* nematode. **b.** The death-associated contraction and expansion of a *P. pacificus* nematode. **c.** The death-associated contraction and expansion of a *C. japonica* male. **d.** A *C. elegans* nematode that was observed to move during the death-associated expansion phase.

**Supplementary Figure 2:**
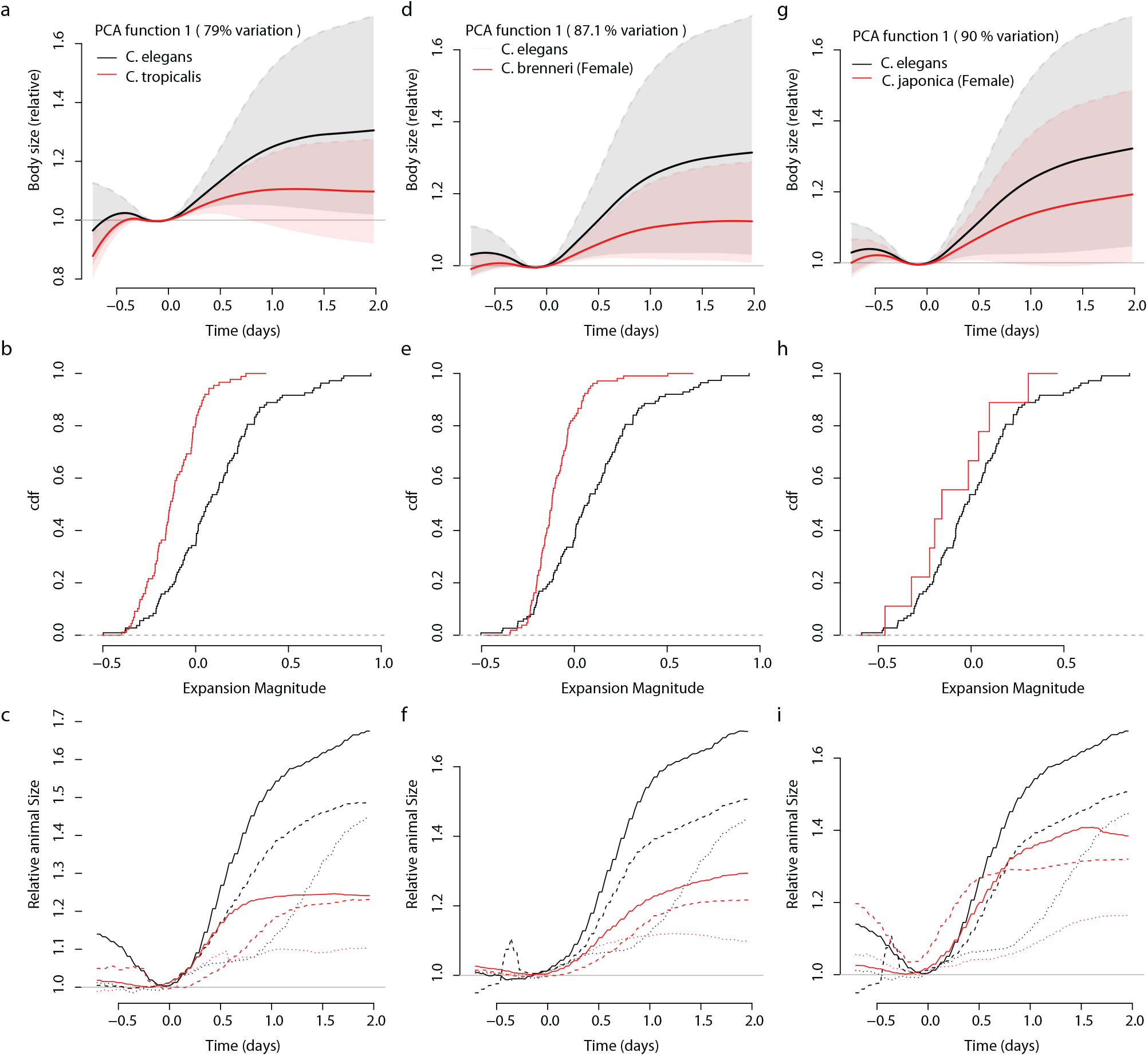
Death-associated morphological changes are conserved across nematode species and paralyzing treatments. **a.** We annotated the movement and morphological transitions by hand for 108 *C. elegans* and 88 *C. tropicalis* nematodes. **b.** The variability in death-associated morphological transitions across both strains was analyzed using FDA, where similar average trajectories (solid line) as well as a similar inter-individual variability (+1 standard deviation: shaded region). The first principle component value of the functional PCA allows us to compare the magnitude of death-associated morphological change between individuals and populations. **c.** To place the PCA in context, we show an example trajectory of animals exhibiting a median magnitude death-time expansion *(dashed line)*, low-magnitude death-time expansion (dotted line), and an high-magnitude death-associated expansion *(solid line)*. We performed a similar analysis of death-associated expansion comparing **(d-f)**, 113 *C. elegans* to 105 *C. brenneri* nematodes, **(g-i)**, 113 *C. elegans* to 9 *C. japonica* nematodes.

**Supplementary Figure 3:**
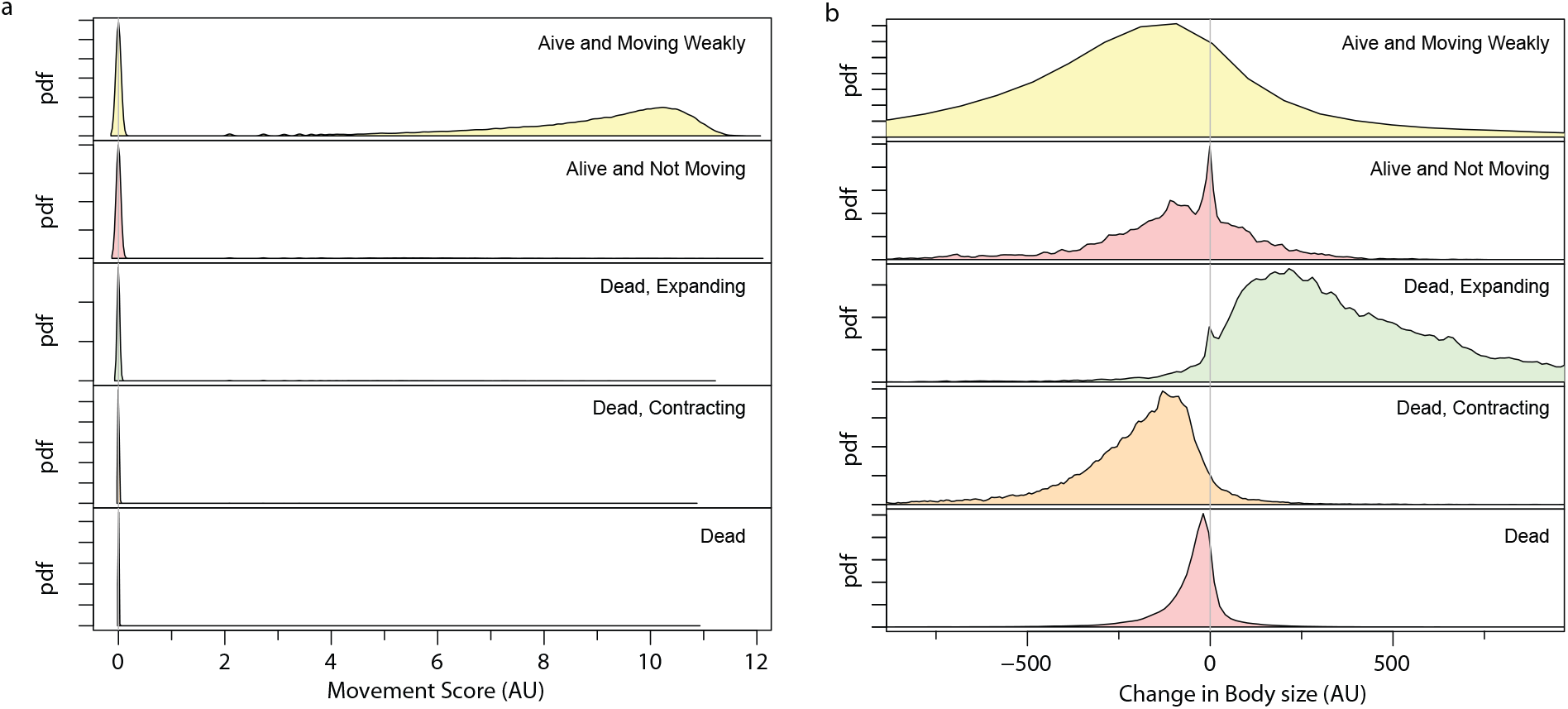
Death-associated morphological changes support robust automated identification of nematode death times. **a.** In an manually-curated data set of 351 individuals, we calculated the empirical probability distribution function for all observations of each animal in each state. **b.** A empirical probability distribution functions quantifying the magnitude of change in apparent body size observed for each animal in each state.

**Supplementary Figure 4:**
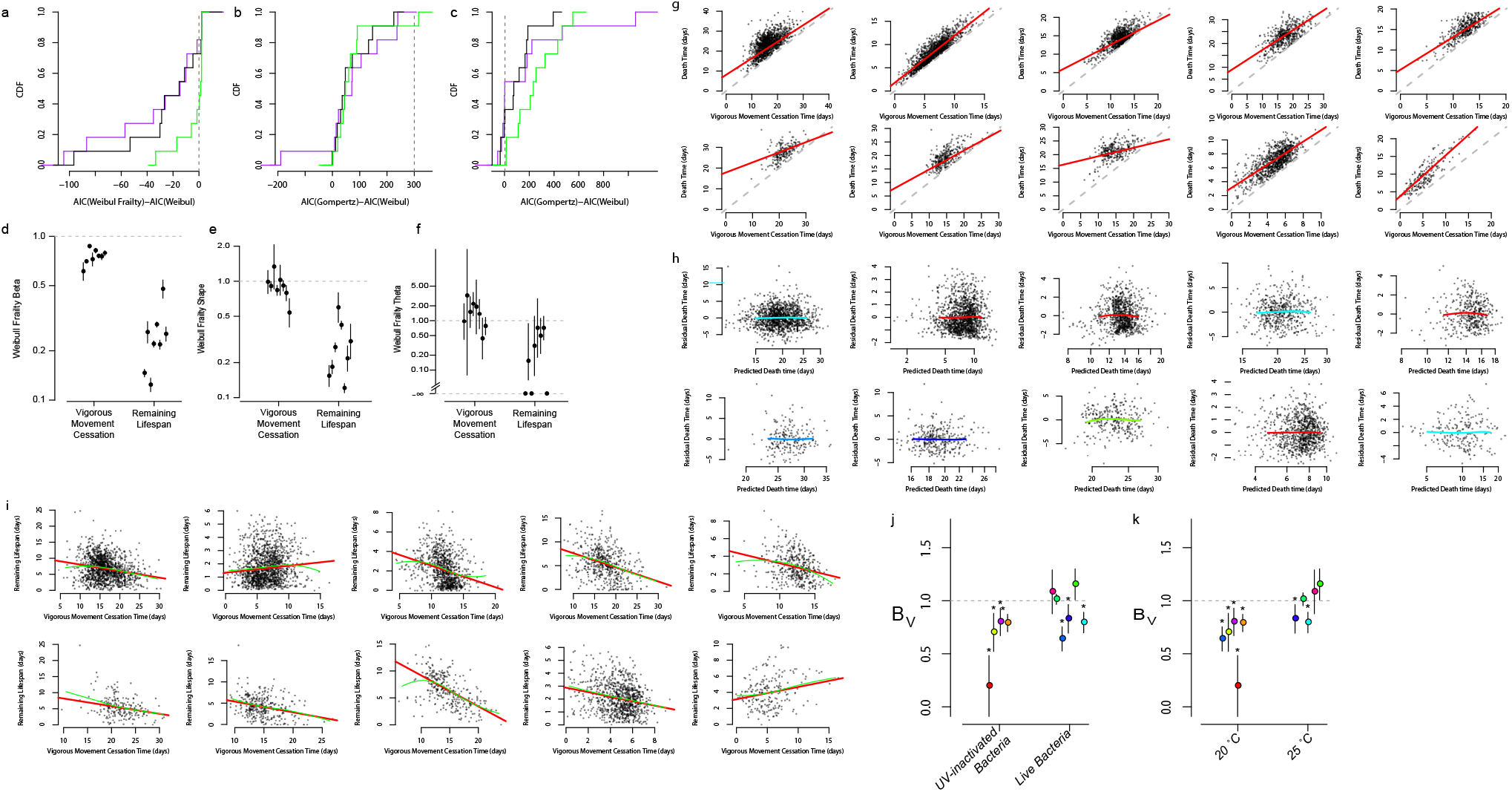
Relating behavioral stages and lifespan in wild-type *C. elegans*. **a.** The cumulative distribution function (CDF) across ten replicates, showing the difference between the AIC criterion obtained from the three parameter Weibull Frailty model, compared to the two parameter Weibull model. Purple is the CDF for models fit to vigorous movement cessation times, black is the CDF for models fit to death times, and in green is the model fit to the remaining lifespan after vigorous movement cessation. Negative values indicate a preference for the three parameter Weibull over the two parameter model. **b.** The same model selection procedure, but this time comparing the Inverse Gaussian distribution to the two parameter Weibull model. **c.** The same model selection procedure, but this time comparing the Gompertz distribution to the two-parameter Weibull distribution. **d.** A parametric regression was performed comparing three-parameter Weibull fits of death times, vigorous movement cessation times, and the distribution of remaining lifespan after vigorous movement cessation. Plotted are the coefficients that relate the scale parameter (beta) for vigorous movement and remaining lifespan distributions to the same parameter of death time distributions. The scale factors all all negative, reflecting the constraint that vigorous movement and remaining lifespan must always be less than total lifespan. **e.** The same regression results, but this time comparing the three-parameter Weibull shape (alpha) parameter. **f.** The same analysis, but this time comparing the Weibull frailty parameter (theta). **g.** The linear regression analysis performed in Fig 4. g-j was repeated for each replicate individually. *h.* The residuals are plotted from the linear regressions in g., with LOESS regression lines overlaid. **i** The relationship between vigorous movement and remaining lifespan, shown in main text Fig. 4e, are plotted for each biological replicate. Linear regression lines *(red)* are compared to the LOESS regression line *(green)*.**j.** In each replicate, the slope of the linear regression line was calculated and plotted. Slopes are grouped according to the food source of the population being described. **k.** The same regression slopes, but grouped according to the temperature at which animals were housed. Because most conditions were performed either on UV-inactivated bacteria at 20 °C or on live bacteria at 25 °C (these being the two standard conditions in the lab), one must refer to Supplementary Figure 5 to differentiate between the influence of food source and temperature. Panels j and k merely indicate that negative slopes were observed with both food sources and at both temperatures.

**Supplementary Figure 5:**
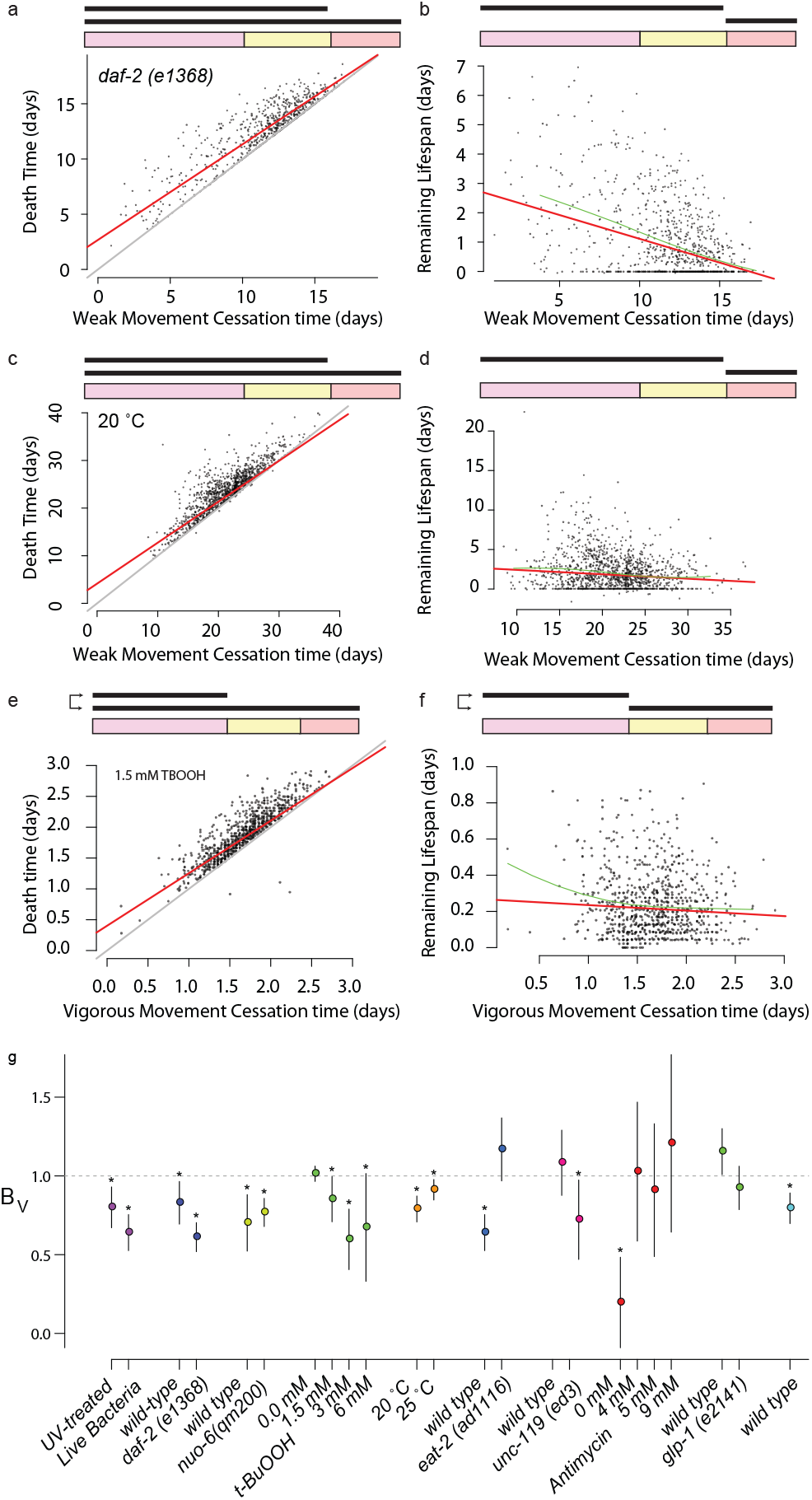
The effect of interventions on behavioral stages and lifespan. a-f. We considered the same data and analyses presented in the main text Fig. g-j, but here show the relationship between the timing of cessation of weak movement and lifespan for a-b *daf-2(e1368)* animals and c-d individuals housed at 20 °. e-f We replicate the analysis of main text Fig. 5 k-l comparing the vigorous movement and lifespan of animals, but this time for animals on 1.5 mM t-BuOOH. a. For all populations considered in Figure 7, we quantified the correlation between the time at which individuals ceased vigorous movement and their lifespan, summarized by the estimate *β_V_* (statistical methods). Significant deviations from *β_V_* = 1 were estimated by bootstrapping and marked with a *. b. For just the wild-type populations from panel a, we compared the *β_V_* of all groups based on whether the experiment was performed on uv-inactivated or live bacteria, and also c. whether the experiment was performed at 20 °C or 25 °C. Because most conditions were performed either on UV-inactivated bacteria at 20 °C or on live bacteria at 25 °C, these being the two standard conditions in the lab, we can only infer from panel a that the differences between groups are primarily due to the temperature (which increases *β_V_*) and not bacteria (which decreases *β_V_*). However, panels b. and c. highlight that significant deviations of *β_V_* below 1 can be observed in all conditions. For all populations considered in Figure 5 and 6, we estimated *β_V_* , the linear regression slope relating vigorous movement to lifepsan, summarized by the estimate (statistical methods). Significant deviations from *β_V_* = 1 were estimated by bootstrapping and marked with an asterisk.

**Supplementary Figure 7:**
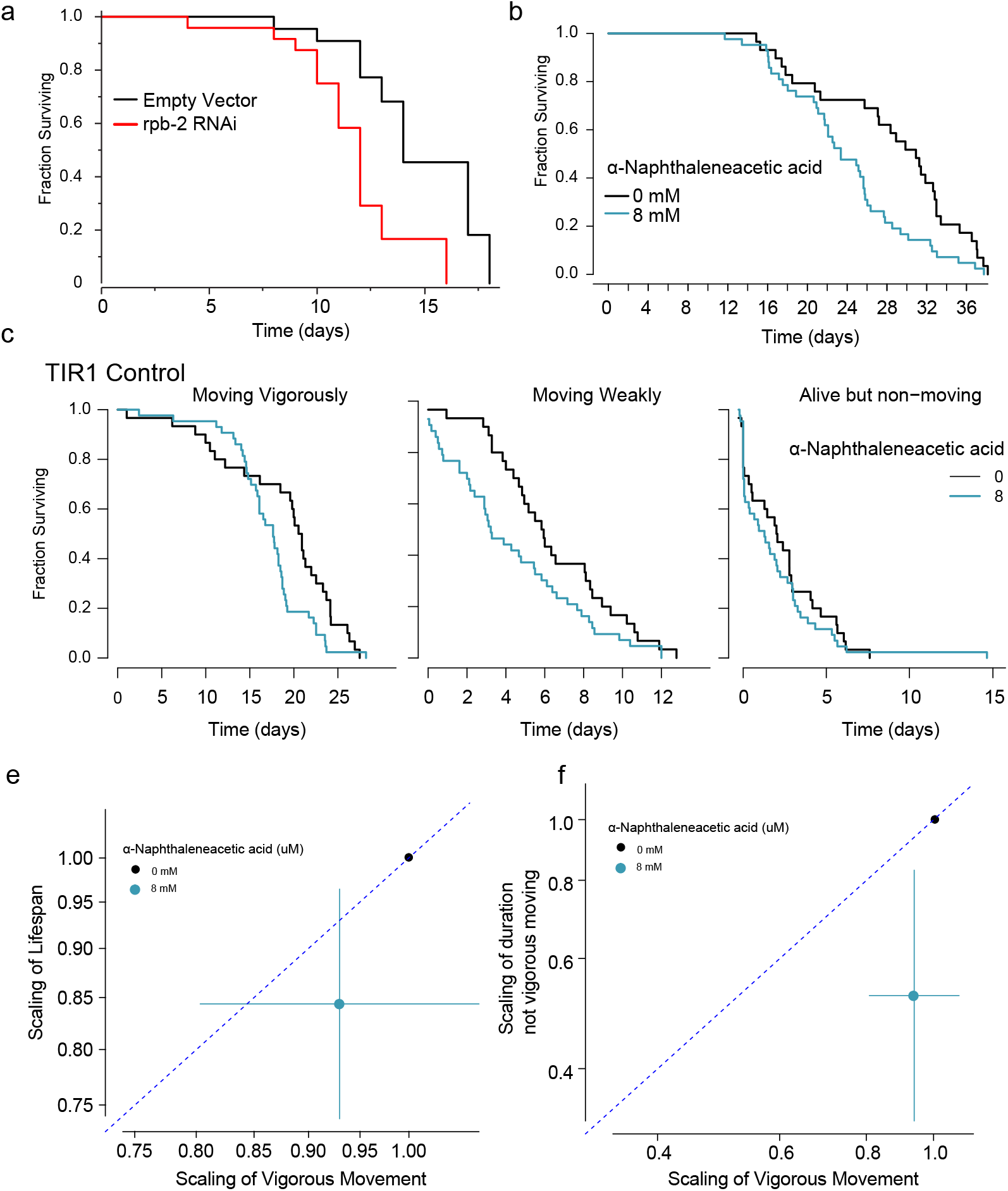
An RNA Polymerase II dosage series. a. On the second day of adulthood, animals living on HT115 at 25 C were transferred from empty vector to rpb-2 RNAi and the Kaplan-Meier survival curve was estimated. c. The durations spent moving vigorously, weakly, and non-moving were compared for TIR1 animals on 0 and 8 mM alpha-Naphthaleneacetic acid (NA). d. The durations spent moving vigorously, weakly, and non-moving were compared for rpb2::AID ; TIR1 animals on 0 and 8 mM NA. e. The relative effect of 8mM KNA on the vigorous movement (93% (80% - 107%); p= 0.33) and lifespan (84% (73% – 96%); p=0.01) was estimated for TIR1 animals lacking the rpb-2::AID degron tag. f. The relative effect of 8mM KNA on the vigorous movement (93% (80% - 107%); p= 0.33) and the duration spent not moving vigorously (52% (33% - 83%); p = 0.006) was estimated for TIR1 animals lacking the rpb-2::AID degron tag. f. The relative effect of KNA dosage on the time spent moving vigorously and the time spent not-moving vigorously was estimated for rpb2::AID ; TIR1 animals.

**Supplementary Figure 8:**
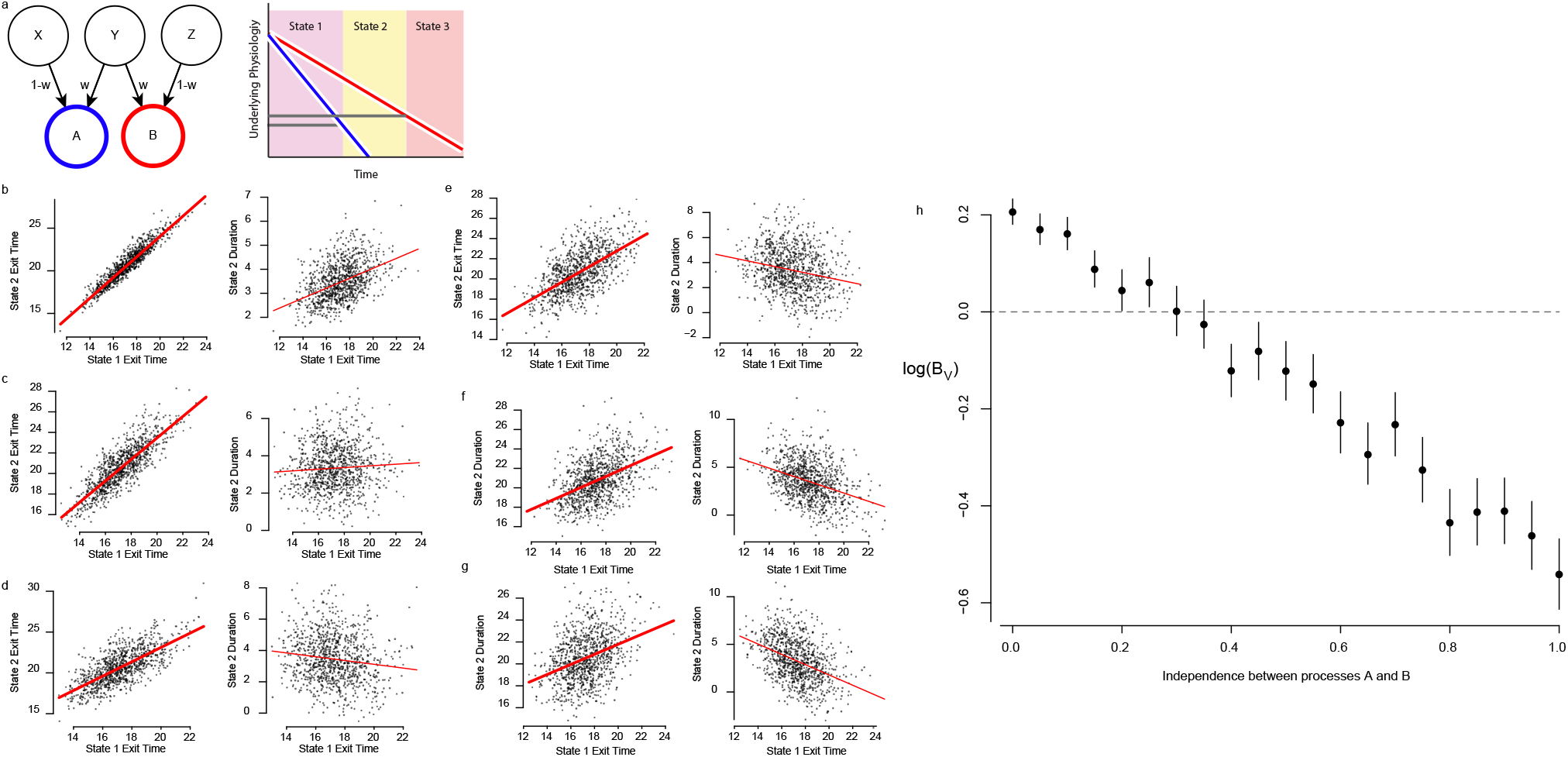
Stochastic model link physical decline to state transition times. . To test our ability to infer dependencies among biological processes using only state transition times, we articulated a model that allows us to vary the dependence between two biased random walks (Weiner processes). a. In this model, the step size of two Weiner process A *(red)* and B *(blue)* at each step of the simulation are determined as the weighted sum of random variables X and Y, and the weighted sum of Y, and Z respectively. As in Fig. 6e, the drift term of both processes vary between individuals to simulate frailty. By changing the weight w, the relative contribution of the independent components (X, Z) and the shared component (Y) can be adjusted. b. When w=1, the two Weiner processes depend only on the shared component Y, and therefore produce a positive correlation between state exit times *(left)* and a positive correlation between state 1 exit time and the duration of state 2 *(middle)*. When the population is stratified by the timing of entry into state 2, individuals who enter later tend to remain longer in state 2 *(right)*. c. At w=0.2, the increased independence reduces the correlation decreases the correlation between state entry times (left) and between state 1 entry and state 2 duration. d. The correlation further decreases when w=0.4, e. w=0.6, f. w=0.8, and g. w=1.0. At w=1.0, the two processes are completely independent and the simulation is equivalent to main text Fig. 8e. h. We then plotted the slope of the regression lines *(red lines, panels b-g)* as a function of w.

## 1 Statistical Methods

### Functional Data Analysis (Figs 1–2)

Functional data analysis was performed by fitting each individuals’ cropped time series with a third-degree basis spline with eight knots, and then performing a PCA on the resulting spline fits. FDA analysis was performed in R using the package fda. Time series describing changes in late-life body size were obtained using the lifespan machine worm browser, and cropped to focus on an interval starting three days prior to and four days after each individuals’ death.

### State-specific score PDFs (Figs 3, Sup. Fig 3)

By hand-annotated movement scores and change in body size measurements were obtained using the lifespan machine worm browser. PDF estimates were obtained using a Gaussian kernel density estimator in R.

### Hidden Markov Model Estimation and Evaluation (Fig. 3)

Hidden Markov Models, parameter estimation, and cross-validation routines were implemented in C++ and integrated into the lifespan machine github repository. In each round of cross validation, the movement scores and body size change measurements present in that round’s training sets were fit with Gaussian Mixed Models (GMMs) and exponential distributions, respectively. GMMs were fit using the GMM library https://github.com/luxiaoxun/KMeans-GMM-HMM . These GMM and exponential fits were then as parameters to our implementation of the Viterbi algorithm, based on the Press, Teukolsky, Vetterling, and Flannery. Numerical Recipes: the art of Scientific Computing Third Edition.

### Transition-Specific Hazard Functions (Fig. 4–5)

The risk of state transitions was calculated as a function of chronological age through numerical differentiation of the Kaplan-Meier cumulative hazard estimate in R. Clock-reset hazard models were calculated by subtracting vigorous movement cessation times from each individuals’ lifespan and numerically differentiating the cumulative hazard estimate of that set in R **Parametric fits (Supp. Fig. 4)—** Parametric fits were estimated using the following forms:

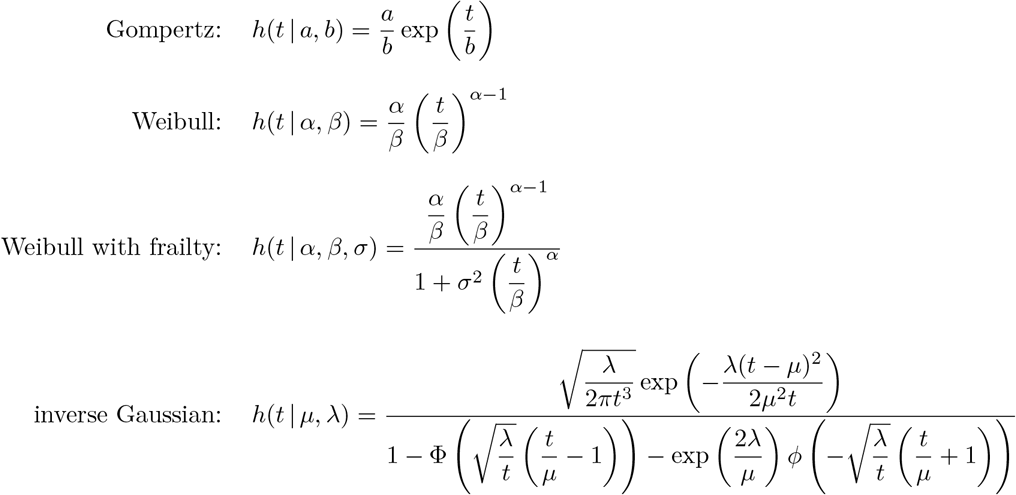

A more in-depth description of these parameterizations is available in the Supplementary information of Stroustrup N, Anthony WE, Nash ZM, Gowda V, Gomez A, López-Moyado IF, Apfeld J, Fontana W. The temporal scaling of Caenorhabditis elegans ageing. Nature. 2016. All parametric fits, along with parametric regression models, were performed using the flexsurv package in R.

### Comparison between state transition times (Fig 4–5,8)

The relationship between state transition times was explored using the multiple regression model with which we control for the confounding influence of batch effects:

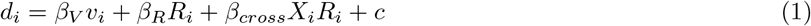

where *V_i_* is the time of cessation of vigorous movement and *d_i_* is the death time of individual *i*. *R_i_* is a categorical variable coding for the batch of individual *i* which in all cases represented the specific flatbed scanner on which the individual was observed. A variety of residual forms and link functions were explored using a generalized linear model approach. We found that standard linear regression provided the best performance, and so the regression was solved using rlm using the ‘RMS’ package in R. Partial coefficients of determination were calculated by comparing the full model with a reduced model with the reduced model *y_i_* = *β_R_R_i_* + *α* using the r package ‘rsq’. A more in depth exploration of this regression approach is presented in Supplementary Notes 1 and 2. The same model was run to evaluate weak movement span, with *v_i_* then representing the timing of cessation of weak movement. LOESS regression was performed in R.

### Accelerated Failure Time models (Fig 6–7)

Scale factors were estimated for each intervention, estimating its effect on vigorous movement cessation time, weak movement cessation time, and death times separately. In each case the following model was fit:

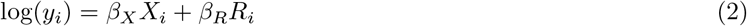

where *y_i_* is the event time under consideration, *X_i_* is a categorical dummy variable coding for the genotype or environmental condition being evaluated, and *R_i_* is the batch effect (in this case the flatbed scanner on which the individual was housed).

### Stochastic process simulations (Fig 6, Supp. Fig 6)

We simulated a one dimensional Weiner processes with drift that satisfies the property that between times *t*_1_ and *t*_2_, individuals advance in their positions *W* (*t*) such that that

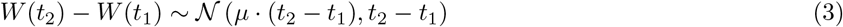

where 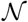 is the normal distribution representing a drift of *μ* relative to the variance *t*_2_ − *t*_1_. Each simulation considered the progression of 1000 random walkers, starting at a displacement from the origin of about 4*x*10^5^ times larger than the drift term. Time was segmented into even steps, and in each step each walker advanced towards zero according to the above *W* (*t*_2_) − *W*(*t*_1_) property. The state transition time of each individual was identified as the first time step in which a walker’s position fell below a constant threshold. For single process models, sequential state transitions correspond to walkers’ passing sequentially lower thresholds. For process models involving frailty, each random walker was assigned a characteristic drift rate, distributed according to a gamma distribution with equal shape and scale parameters, centered around the population average drift-rate *μ*. For multiple process models, walkers progressed according two independent Weiner processes, each with its own drift *μ* and threshold governing state transition. For the partially independent processes in Fig 6, two coupled Weiner processes *W_A_*(*t*) and *W_B_*(*t*) were considered that satisfy the property that

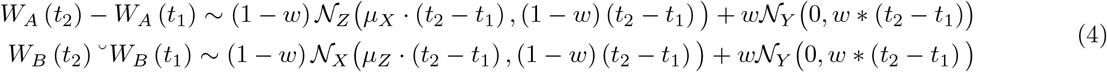

where 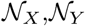 ,and 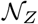 are independent variables. Because of the property of normally-distributed variables that 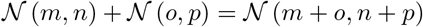, in our simulation changing the constant w will not alter the characteristic drift or random components of the two Weiner processes *W_A_* and *W_B_*, but instead will change only the magnitude of dependence that their advancement at each simulation step has on the shared 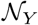 variable. As in previous multiple process simulations, state transition times are identified as the first step in which *W_A_*(*t*) and *W_B_*(*t*) pass below their respective thresholds.

