## Supplemental Notes 1 and 2 for "*A Hierarchical Process Model Links Behavioral Aging and Lifespan in* C. elegans"

#### 1 Supplementary Note 1–relating correlations, slopes, and processes

We assume there are at least two underlying sources of biological variation in our data set. The first is $V$ which is the endpoint distribution of some underlying stochastic process determining cessation of vigorous movement. There are two additional sources that we are unsure about, the first being $D$ which is another endpoint distribution describing death times. The second is $X$, which describes how much longer an individual lives after it ceases moving vigorously, such that if for each individual $i$, we measure their event times $d_i \sim D$ and $v_i \sim V$, then $x_i = d_i - v_i$ with $x_i \sim X$. For all individuals, we observed that $d_i \geq v_i$, with $E(D) \gg E(V)$. Our goal in this document is to understand how our pairwise data $(v_i, d_i)$ can be used to understand the relationship between $V$, $D$, and $X$.

One possible relationship between $V$, $X$, and $D$ is shown in Fig. 1

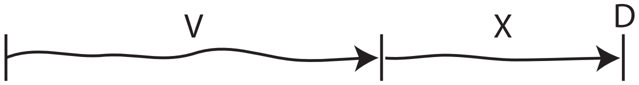


**Figure 1:** A model where $V$ and $X$ run in series

In this model, each death time $d_i$ occurs at the end of a process $X$. For each individual $i$, process $X$ begins at time $t = v_i$ generating the relationship $d_i = v_i + x_i$. In other words: first individuals cease vigorous movement and then they start dying.

This model could reflect an interplay between qualitatively distinct physical phenomena determining $D$ and $V$. Cessation of vigorous movement could trigger a new process that ultimately kills the animal. However, this model also describes the case where $D$ and $V$ represent manifestations of a single underlying process. For example, $V$ and $D$ might reflect two first passage times of a single random walk, with $v_i$ and $d_i$ being first passage time across two distance thresholds. In the case of random walks specifically and more generally among memory-less stochastic phenomena, $D$ and $X$ would be independent even though $D$ and $V$ are correlated. This results from essential sequential nature of this model, in that $D$ is correlated because $d_i = v_i + x_i$. The independence of the generative processes $X$ and $V$ becomes obvious by conditioning $D$ on $V$.

In our empiric data, we relate $D$ and $V$ via linear regression, and find the slope is less than one. To put this in context of our model here, we start with the case where $D$ and $X$ are independent. Therefore, $E(D) = E(V) + E(X)$, for all times $t$ and so

$$E(D|V = t) = t + E(X|V = t) = t + E(X) \tag{1}$$

When we run the regression

$$d_i = \alpha v_i + \epsilon_i + c \tag{2}$$

on our paired data $(d_i, v_i)$, we obtain the estimate of $\alpha$ that ensures that

$$E(D|V = t) = \alpha t + c \tag{3}$$

with $E(\epsilon|V*) = 0$ Eq. 1 ensures that model (1) when $X$ and $D$ are independent will generate data for which estimates of $\alpha$ will always be 1. In contrast, in our empiric data suggests that $\alpha \leq 1$.

We can alter model (1) to allow for $\alpha < 1$ only by introducing a negative correlation between $V$ and $X$. This can be shown by starting at the relationship $D = V + X$, which implies that $E(D|V = t) = E(D|D - X = t) = E(D|D = t + X) = t + E(X|V = t)$. Because regression 2 solves for 3, we can say that

$$E(X|V = t) = (\alpha - 1)t - c \tag{4}$$

Therefore, when $\alpha$ is less than one, this implies that $E(X|V = t) = \gamma t - c$ with $\gamma < 1$. This can only arise when $X$ and $V$ are negatively correlated.

So, how could such a negative correlation arise between $X$ and $D$? To investigate this, we must be more explicit about the causal relationship between $V$ and $D$. Consider two causal relationships between $V$, $D$, and a potential upstream confounder $R$.

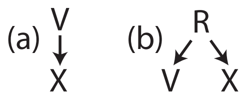


**Figure 2:** Two possible causal relationships between $V$ and $X$

Causal models (a) and (b) are distinguished by the relationship between $V$ and $D$, which in (a) is causal and in (b) is not causal. In (a) $\alpha < 1$ implies that $V$ exert a negative influence on $X$. Individuals with large a $d_i$ will have a small $x_i$ *because* they have a large $v_i$, compared to the population average. In other words, remaining vigorous for longer must somehow cause animals to subsequently live shorter.
In contrast, model (b) allows the correlation between $V$ and $D$ to arise from the action of some upstream factor $R$. Because $V$ is no longer causally upstream of $V$, then the inverse correlation implied by Eq. 4 need no longer reflect a physical interaction between $V$ and $D$ that persists after $t = v_i$. Instead, the inverse correlation between $X$ and $V$ and negative $\alpha$ would arise as a statistical epiphenomenon determined by whatever arbitrary form $E(D|V = t)$ results from the relationship between $E(D|R = i)$ and $E(V|R - i)$. Consider the two extreme cases of model (b): The case where $R$ fully determines $V$ and $X$, at which point $\alpha = 1$, and the case where $R$ has no effect on $V$ and $X$, at which point $\alpha = 0$. Any realization of model (b) that falls these extreme cases must then produce an $\alpha$ whose value lies between 0 and 1.

These constrains also hold if we allow $V$ and $D$ to progress in parallel, as shown in Fig. 3

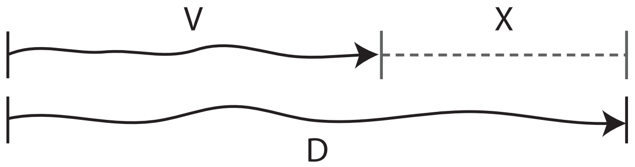


**Figure 3:** A model where $V$ and $D$ run in parallel

Here, $d_i \geq v_i$ remains true only because $E(D) \gg E(V)$ and perhaps as the result of of rare truncation events where $D$ truncates $V$. Causal model (a) requires that the process determining $V$ be coupled to $D$ such that $D$ progresses slower before $t = v_i$ but then faster after $t = d_i$. In contrast, causal model (b) allows $V$ and $D$ to progress without direct interactions, at rates similarly influenced by the upstream factor $R$. We therefore conclude that estimates of $\alpha$ in Eq. 2 provide a means for differentiating among possible relationships in the physiologic determinants of $V$, $D$ and $X$. $V$ and $D$ cannot be the outcome of two sequential, independent processes, such as a random walk. Instead, it's possible either that $V$ causally influences $D$ in a way that individuals remaining vigorous for longer subsequently die sooner. Alternatively, if we allow for an upstream factor that can influence both $D$ and $V$, then the observed $\alpha < 1$ can arise in the absence of any causal interactions between $V$ and $X$.

### 2 Supplementary Note 2–Accounting for environmental effects and measurement error

Unfortunately, we cannot directly observe $D$ or $V$. Our observations are distorted by the confounding effects environmental effects that we summarize as a single factor $Z$ as well as measurement error $\epsilon$. We can only observe $D*$ and $V*$:

$$\begin{aligned} v_i^* &= v_i + z_{v_i} + \epsilon_{v,i} + c_v \\ d_i^* &= d_i + z_{d_i} + \epsilon_{d,i} + c_d \end{aligned} \tag{5}$$

with $E(\epsilon_{v,i}) = E(\epsilon_{d,i}) = E(z_{v,i}) = E(z_{d,i}) = 0$ and $Var(\epsilon_{v,i}) = \sigma^2_{ev}$, $Var(\epsilon_{d,i}) = \sigma^2_{ed}$, $Var(z_{v,i}) = \sigma^2_{zv}$, and $Var(z_{d,i}) = \sigma^2_{zd}$. In this application, there can be no measurement error in $Z$, as $Z$ is a categorical variable coding for a discrete organizational unit (the plate or device on which the measurements were obtained) predefined before the start of the study.
In regression 2, we seek to estimate the relationship $d_i = \alpha V + \epsilon_{b,i}$. Combining this with 5, we find that the relationship between observed variables $D*$ and $V*$ is

$$d_i^* = \alpha v_i^* - z_{d_i} + \alpha z_{v.i} + \epsilon_{b,i} - \epsilon_{d,i} + \alpha \epsilon_{v,i} - c_d + \alpha * c_v \tag{6}$$

The correlation between $v_i$, $Z$, and $\epsilon_{v_i}$ is an unfortunate example measurement error's effect on regression. We wish to estimate $\alpha$ in the presence of environmental factors $Z$ using the multiple regression model

$$D_i = \beta_V V_i + \beta_Z X_i + \epsilon_i; \tag{7}$$

However, the influence of $\alpha$ on $V$, $Z$ and $\epsilon$ in Eq. 6 introduces structural problems in such estimation. Multiple regression in the presence of measurement error is well studied (Buonaccorsi. Measurement Error: Models, Methods, and Applications) (E. S. Gillespie. The influence of measurement errors in multiple linear regression.), and predicts that $\beta_V$ can be a biased estimator of $\alpha$. The nature of this bias will depend on the correlation between $Z$ and $V$, as well as the relative magnitudes of $\sigma^2_{ev}$, $\sigma^2_{ed}$, $\sigma^2_{zv}$ and $\sigma^2_{zd}$. To avoid extended analytic excursions, in the following sections we follow two approaches to identify any bias in our estimation of $\alpha$ using regression 7.

#### 2.1 Bias in multiple regression estimates I : Estimating $\alpha$ within batches

First, we study the relationship between our estimates of $\alpha$ across the whole data set to estimates obtained in subsets of the data. We consider the data presented in Fig. 5 of the main text which describes 2346 individuals, organized into four batches. Each batch represents a flatbed scanner, which produces characteristic set of unmeasured environmental factors (e.g temperature) that influence the $V$ and $D$ of all animals housed on that scanner. As previously shown elsewhere, regression 7 estimates $\alpha$ at a value significantly below 1. Without accounting for batch effects, the single regression model 2 fails to identify a significant deviation from $\alpha = 1$, highlighting the importance of accounting for batch effects. To see whether $B_Y$ in the multiple regression 7 is a biased estimator of $\alpha$, we considered the relationship between $D$ and $V$ in each batch separately, using the single regression model 2. In table 1, we find that when grouped with each batch-estimated $\beta_y$, the all-batch estimate of $\beta_y$ is the median value. This suggests that the structure of our multiple regression approach (Eq. 6) does not add substantial bias to our estimate of $\alpha$ on top of what is present in a single variate regression model of the same data.

| Data used | Model | N | $\beta_Y$ | 95% CI | p-value (a <1) | Intercept | 95% CI |
|---|---|---|---|---|---|---|---|
| All | $d_i = \beta_V V_i + \beta_Z Z_i$ | 1380 | 0.86 | (0.82-0.91) | <.0001 | 4.83 | (4.23-5.43) |
| All | $d_i \beta_V V_i$ | 1380 | 0.98 | (0.93-1.02) | 0.1432 | 3.49 | (2.93-4.05) |
| Batch 1 | $d_i \beta_V V_i$ | 312 | 0.82 | (0.72-0.92) | 0.0006 | 6.00 | (4.73-7.26) |
| Batch 2 | $d_i \beta_V V_i$ | 430 | 0.78 | (0.70-0.87) | <.0001 | 6.33 | (5.24-7.42) |
| Batch 3 | $d_i \beta_V V_i$ | 391 | 0.97 | (0.89-1.05) | 0.2666 | 2.49 | (1.63-3.35) |
| Batch 4 | $d_i \beta_V V_i$ | 247 | 0.89 | (0.79-0.98) | 0.0054 | 4.56 | (3.36-5.77) |

**Table 1:** Comparing $\alpha$ estimates within and across batches

The values of $\beta_D$ in table 1 suggests that our multiple regression approach does not introduce additional bias compared to single regression model estimates. The estimate of $\beta_v$ obtained via multiple regression is centered

among batch-specific $\beta_v$ estimates.
It still remains the case that these single regression model estimates may themselves be biased due to the measurement error term $\epsilon_{v,i}$ in Eq. 5. Measurement error in single regression produces attenuation in parameter estimates that case might make $|\beta_D| \leq |\alpha|$. This would impede our ability to distinguish between causal models (1), (2), and (3) by preventing us from distinguishing between $\alpha = 1$ or $\alpha < 1$. Fortunately in this case analytic theory is more straightforward than for multiple regression. Attenuation will take the form $|\beta_V| = |\hat{\kappa}\alpha|$ with

$$\hat{\kappa} = \frac{\hat{\sigma}_D^2}{\hat{\sigma}_D^2 + \hat{\sigma}_{Derr}^2} \tag{8}$$

Here, $\hat{\sigma}_V^2$ and $\hat{\sigma}_{Verr}^2$ are the variances of $V$ and the measurement error in $V$ in eq. 5. By substituting in our empiric estimate $\beta_V = 0.8$ and asserting $\alpha = 1$, we can use eq. 8 to predict the relative magnitude of measurement error required to produce a bias large enough to generate the observed deviation from $\beta_v = 1$. We find that $\hat{\sigma}_{Verr}^2 = 0.25\hat{\sigma}_V^2$, so that a measurement error with a standard deviation equal to half of $V$ would be required produce the observed bias. Unfortunately, the nature of our time-to-event data is that repeated measurements are impossible, as each individual's cessation of vigorous movement and death can only be observed once. It's therefore impossible to directly estimate the true empiric ratio of $\hat{\sigma}_{Verr}^2/\hat{\sigma}_V^2$. However, we can inspect various aspects of our data to estimate the contribution of measurement error to our analysis.
In the wild-type data set shown in main text Fig. 5, $\hat{\sigma}_V = 2.2$. Numerical simulation using a Weibull distribution $V$ and additive Gaussian noise for measurement error, we found that the underlying biological noise and measurement error would combine to show $\hat{\sigma}_V = 2.2$ when $\sigma_V = 1.99$ and $\sigma_{Verr} = 1.00$, as shown below.

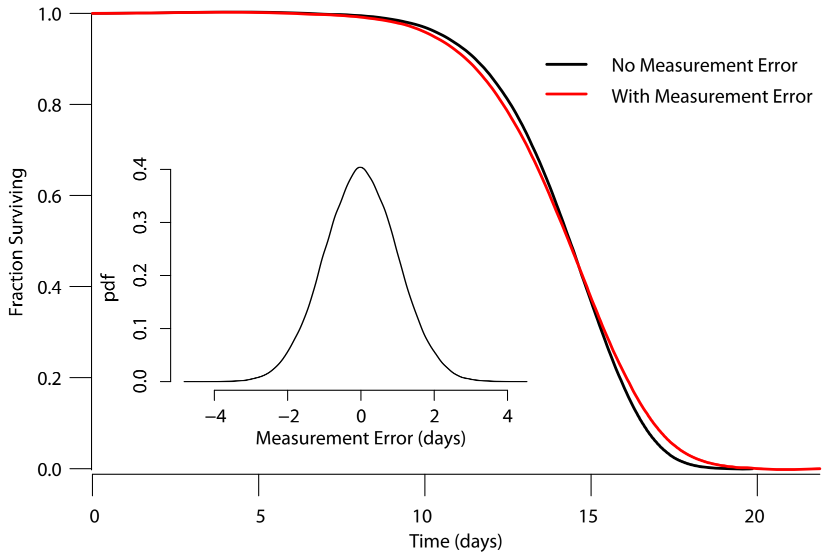


**Figure 4:** Additive error with the standard deviation such that $\hat{\sigma}_{Verr}^2 = 0.25\hat{\sigma}_V^2$ is shown in the PDF inset. This error has the effect of transforming the "true" distribution (black) into an "observed" distribution (red).

Fig. 4 provides an example in which measurements of an individuals' vigorous movement span would need to have an average absolute error of 0.8 days. This is roughly two-fold higher than the typical error of our automated analysis pipeline, and subsequent by-hand annotation of our data sets reduces the error to less than 0.1 days error on average. Yet, reducing error below 0.8 days on average is a taxing standard, and so we performed additional diagnostics to see whether the observed $\hat{\sigma_V}$ or other population level statistics correlated with the observed $\beta_V$.

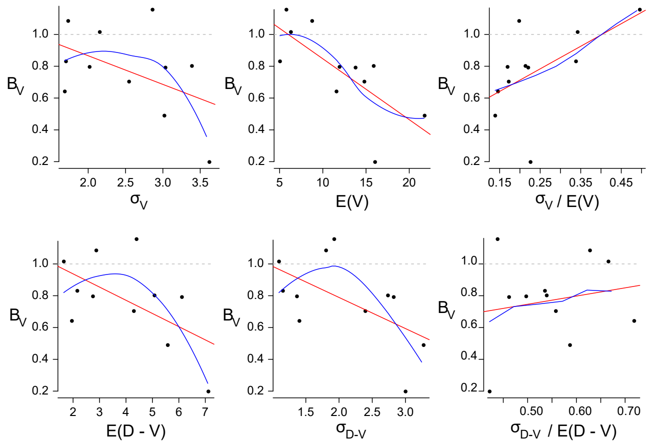


**Figure 5:** The relationship between observed $B_v$ and population summary statistics across replicate experiments

We found that the $\beta_v$ was inversely correlated with $\sigma_V$, but also with the $E(V)$. Were $V$ to vary across replicates by a scaling relationship, we'd expect proportional changes in $\sigma_V$ and $E(V)$. So, we calculated the coefficient of variation to understand the "mean-normalized" with our $\beta_v$ estimate. We found that "less noisy" experiments in fact showed lower $\beta_v$, not supporting the hypothesis that measurement-error is attenuating our parameter estimates. We then considered the relationship between $D$ and $V$ and $\beta_V$. Larger intervals $v_i$ $d_i$ correlated with smaller $B_v$, but so did the standard error $\hat{\sigma}^2_{D-V} = var(d_i - v_i)$. So, we calculated the coefficient of variation for $D - V$, and found no relationship with $B_v$ estimates. In conclusion, though we identified relationships between $\beta_V$ and various population statistics including $\sigma_V$, accounting for the understood mean-variance relationship between distributions of vigorous movement span, increased variation is related to increased (not attenuated) $\beta_v$ estimates.
Therefore, though we cannot fully exclude the role of measurement error, and eagerly await improvements in measurement accuracy that will definitively resolve the issue, we find no evidence to substantiate theoretic concerns that measurement errors make $\beta_v$ deviate from $\alpha$ in our analysis.

#### 2.2 Simulating the effect of measurement error and environmental effects

To demonstrate our ability to distinguish between causal models (2) and (3) in the presence of environmental noise, we performed a series of simulations.

$$\begin{aligned} &\textbf{Variables shared in both simulations} \\ V^\dagger &\sim Weibull(\beta = 12, \alpha = 5.75) \\ R &\sim Gamma(\beta = 70, \alpha = 1/70) \\ Z &\in \{1, 2, 3, \dots, N\} \\ \beta_Z &\sim Gamma(\beta = 50, \alpha = 1/50) \end{aligned} \tag{9}$$

$$\begin{aligned} &\textbf{Simulation of Causal Model (2)} \\ X^\dagger &\sim Weibull(\beta = 3, \alpha = 2.5) \\ v_i &= r_i * v_i^\dagger \\ x_i &= x_i * x_i^\dagger \\ d_i &= v_i + x_i \\ v_i^* &= v_i * (\beta_Z Z_i) \\ d_i^* &= d_i * (\beta_Z Z_i) \end{aligned} \tag{10}$$

**Simulation of Causal Model (3)**

$$
\begin{aligned}
D^{\dagger} &\sim Weibull(\beta = 15, \alpha = 7.2) \\
v_i &= r_i v^{\dagger} \\
d_i &= r_i d^{\dagger} \\
v_i^{*} &= v_i * \beta_Z Z_i \\
d_i^{*} &= d_i * \beta_Z Z_i
\end{aligned} \tag{11}
$$

The "true" underlying sources of biological variability $V^{\dagger}$, $X^{\dagger}$, and $D^{\dagger}$ are modeled as Weibull distributions with parameters chosen to approximate our empiric data. The causal effect of $R$ is modeled by multiplying each individual's "true" $v_i$ and $x_i$ ( model (2) ) or $d_i$ (model (3)) by a value drawn from a gamma distribution with a mean of 1. $R$ thereby introduces a correlation between $V$ $X$, and $D$ in model (2), and between $V$ and $D$ in model (3). Batch effects, meant to mimic the influence of environmental factors, were introduced by randomly assigning each individual $i$ membership to a batch $Z$. Each batch produces a characteristic effect shared among individuals in that batch. The "observed" $V^*$ and $D^*$ analyzed in the simulation are obtained by multiplying $v_i$ and $d_i$ by the batch effect $\beta_Z Z_i$ characteristic of batch $Z_i$. Note that here we choose to model noise and environmental variation as multiplicative factors in contrast to previous sections, as multiplicative noise is likely to more closely match the physical effect of temperature and other factors on event times (Stroustrup et al 2016.)

First, we performed a simulation in which paired $v_i$ and $d_i$ were generated as samples from the $V$ and $D$ as defined above for each model. The regression model 2 was fit to estimate $\beta_v$, allowing us to evaluate the performance of our statistical approach in the absence of batch effects. Then, we performed the same simulation but this time comparing $v_i^*$ and $d_j^*$ generated as samples from $V^*$ and $^D*$ defined above for each model. The regression model 7 was fit to estimate $\beta_v$, allowing us to evaluate the performance of our statistical approach in the presence of environmental confounders. In both sets of simulations, 1000 replicates were performed each considering a population of 1000 simulated individuals. In addition to estimates of $\beta_v$, we obtained bootstrapped p-values for the test $H_0 : \beta_v < 1$. We summarize these results in table 2, providing the average $\beta_v$ obtained across all replicates as well as the fraction of p-values obtained that fell below a $p = .05$ threshold.

| Data Generating Model | Batch effects added? | $\beta_v$ | Fraction of replicates with significant $\alpha < 1$ at $p < .05$ |
|---|---|---|---|
| Model (2) | No | 0.999 | .044 |
| Model (2) | Yes | 1.000 | .056 |
| Model (3) | No | .520 | 1.000 |
| Model (3) | Yes | .519 | 0.985 |

**Table 2:** Differentiating simulated causal models.

These results show that regardless of environmental effects, we can differentiate between causal models (1) and (2). Causal model (2) generates data matching the expected relationship of $D = \alpha V + \epsilon$ with $\alpha = 1$. Our false positive rate of our bootstrapping test for $\beta_v ! = 1$ matches the expected rate of 5%. Causal model (3) generates data matching the expected relationship $D = \alpha V + \epsilon$ with $\alpha < 1$. The with a false negative rate at this effect magnitude appears at or below 2.5%.

#### 2.3 Power Simulation

To estimate our power to identify $\alpha \neq 1$ and the sample sizes required to detect the deviations we observe, we performed a simulation. Using the data presented in Fig. 5 of the main text which describes 2346 wildtype individuals, we sub-sampled the data set, selecting an equal number of individuals from each batch (device). Across 1000 replicates at each sample size, we then measured the relationship between $\beta_v$ when estimated using a single regression model 2 or the full multiple regression model 6. In each replicate, we calculated the boot-strapped p-value for $\beta_v ! = 1$ for the single and multiple regression models, and estimated the proportion of replicates for which this p-value was significant at $p < .05$

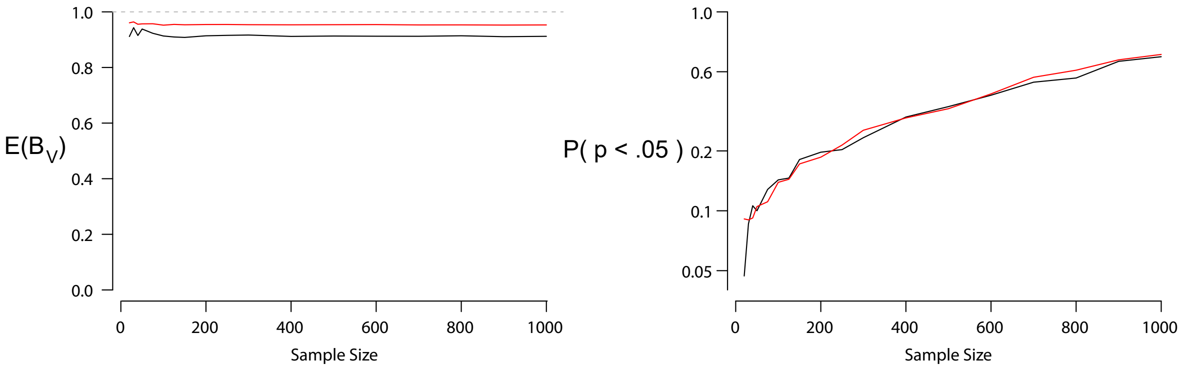


**Figure 6:** Subsampling our empiric data to estimate parameter bias and statistical power as a function of sample size. Multiple regression to account for batch effects (black) and the naive single regression (red).

We found that failure to account for batch effects biased the estimation of $\beta_v$ toward 1 at all population sizes. However, given the small overall magnitude of effect, large populations were required to obtain marginal statistical power, with a power of 0.60 being reached only when 1000 individuals were considered.
